# Pharmacological inhibition of deubiquitinase UCH-L1 by LDN57444 sensitises hepatocellular carcinoma to sorafenib by reverting drug-induced adaptive responses

**DOI:** 10.64898/2026.05.15.725527

**Authors:** Elias Van De Vijver, Kylian Decroix, Delphine Burggraeve, Petra Van Wassenhove, Zenzi De Vos, Christophe Ampe, Lindsey Devisscher, Hans Van Vlierberghe, Marleen Van Troys

**Author notes:** Shared first author (Elias Van De Vijver, Kylian Decroix). Shared last author (Lindsey Devisscher, Hans Van Vlierberghe, Marleen Van Troys). Contact information: Van Vlierberghe Hans Department of Internal Medicine and Pediatrics, Hepatology Research Center, Faculty of Medicine and Health Sciences, Ghent University Campus UZ (1K12 IE), Corneel Heymanslaan 10, 9000 Gent, 09 332 23 70.

## Abstract

**Background and aims:** Therapeutic outcomes for advanced hepatocellular carcinoma remain inadequate, despite recent advances using immunotherapy. Long-term effectiveness of systemic therapies, including second-line multi-tyrosine kinase inhibitor sorafenib, is limited by resistance mechanisms and adverse effects. Upregulated deubiquitinase UCH-L1 is frequently correlated with poor prognosis in cancers. Here, we investigated the therapeutic potential of combining pharmacological UCH-L1-inhibition with sorafenib in HCC.

**Methods:** UCH-L1 expression was analysed in TCGA-LIHC data and patient-derived HCC tissues. Sorafenib and LDN57444 effects were evaluated *in vitro* in cytotoxicity and invasion assays. Gene and protein expression were examined by RT-qPCR, Western blotting and immunohistochemistry. *In vivo* efficacy of drug synergy was assessed in an orthotopic xenograft mouse HCC model.

**Results:** *In silico* data-analysis revealed significantly higher UCH-L1 levels in patient HCC tumours versus non-tumour, associated with reduced overall survival. Low-dose sorafenib upregulated UCH-L1 in HCC cell line Hep3B. Paradoxically, this also promoted invasiveness and sustained MEK1/2-ERK1/2-pathway activation. Combining low-dose sorafenib with LDN57444 produced strong synergistic cytotoxicity *in vitro*, reverted MAPK-activation and suppressed invasion. Consistently, at low sorafenib dose co-treatment with LDN57444 completely inhibited tumour growth of Hep3B xenografts and enhanced sorafenib efficacy.

**Conclusion:** LDN57444 sensitises HCC cells to low-dose sorafenib by reverting drug-induced pro-oncogenic signalling and thereby strongly synergises with sorafenib to enhance anti-tumour efficacy in a HCC mouse model. This presents UCH-L1 as a player in treatment-induced adaptive response and supports further exploring UCH-L1-targeting in combination with sorafenib as therapeutic avenue for advanced HCC.

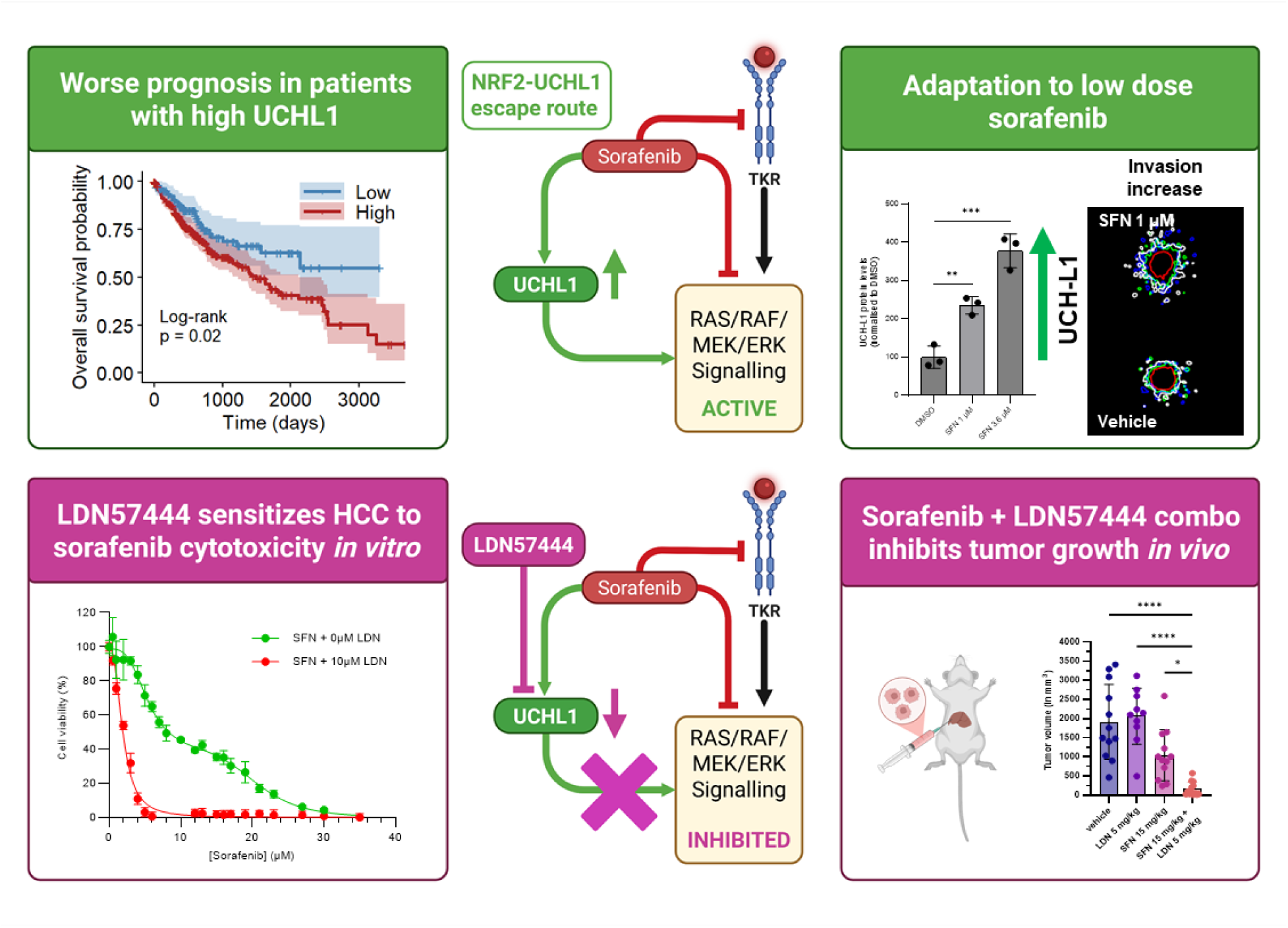

**Lay summary:** This study explores a new treatment approach for hepatocellular carcinoma (HCC) by combining two drugs: LDN57444, which blocks the enzyme UCH-L1, and sorafenib, a FDA-approved multi-tyrosine kinase inhibitor. We evaluated the effect of this drug combination *in vitro* using a HCC cell line and in an mouse HCC-model. The drug combination displayed strong, synergy in lowering HCC cell viability, and greatly reduced invasiveness and *in vivo* tumour growth. LDN57444 sensitised HCC cells to low doses of sorafenib by preventing UCH-L1-mediated activation of pro-oncogenic signalling. These findings highlight the potential of this new drug combination for treating advanced HCC thereby potentially reducing side-effects and countering drug resistance.

**Impact and implications:** Our preclinical research introduces a novel combination strategy against advanced HCC that holds potential to improve existing therapies, particularly the second-line multi-tyrosine kinase inhibitor sorafenib. The proposed combination of sorafenib with an inhibitor of the deubiquitinase UCH-L1 not only enhances sorafenib efficacy but present promise to also counter resistance mechanisms. Moreover, because effective responses are achieved at lower drug doses, this may in addition reduce therapy-associated adverse effects further increasing potential impact. While sorafenib is FDA-approved, the UCH-L1 inhibitor LDN57444 needs further (clinical) development to bring our promising findings to full translational potential for HCC patients and physicians.

## Introduction

Hepatocellular carcinoma (HCC) accounts for over 80% of primary liver cancer cases and is worldwide the third-leading cause of cancer-related mortality^1,2^. The majority of patients are diagnosed at an advanced stage, where therapeutic options are largely limited to systemic therapy. Currently, objective response rates are only 30% and 5-year overall survival rate for advanced HCC (aHCC) is below 30%^1^, underscoring the urgent need for more effective therapies.

Until 2021, multi-tyrosine kinase inhibitors (mTKIs) such as sorafenib were the cornerstones of aHCC treatment^3^. Sorafenib targets multiple membrane receptor kinases (e.g. VEGFR, PDGFR, c-KIT) and cytosolic RAF-kinase, thereby impairing oncogenic RAS-RAF-MEK-ERK signalling^4^. Clinical benefits of mTKIs however have been strongly constrained by substantial adverse effects and rapid occurrence of resistance^5^. The introduction of immune checkpoint inhibitors (ICIs) reshaped the therapeutic landscape of aHCC^6^. Combining atezolizumab (anti-PD-L1) and bevacizumab (anti-VEGF) proved superior over sorafenib, setting it as new standard of care^7,8^. Likewise, combinations using anti-CTLA-4 with anti-PD-L1 or anti-PD-1 proved advantageous over mTKIs^9,10^. Nonetheless, ICI-based therapies still present with low objective response rates and variable primary and acquired resistance, implying they will not suffice to meet clinical needs. Optimaltreatments after failure to ICI-based therapy moreover are also still underexplored.

mTKI-based therapy therefore remains a viable option, particularly within combination strategies. The phase-III CARES-310 study (mTKI rivoceranib plus anti-PD-1 camrelizumab) e.g. demonstrated significantly improved progression-free and overall survival compared to sorafenib^11^. Ongoing studies investigating sorafenib-based combinations with other targeted therapeutics, e.g. mTOR-inhibitor everolimus^12^ and PI3K-inhibitor copanlisib^13^, further highlight future opportunities for innovative mTKI-based combination regimens.

In this study, we propose targeting the deubiquitinase (DUB) UCH-L1 as novel strategy to enhance mTKI efficacy. DUB enzymes are molecular all-rounders, playing essential roles in regulating protein homeostasis and cellular functioning and cancer^14,15^. DUB UCH-L1 is a ubiquitin C-terminal hydrolase^16,17^, displays predominant expression in brain and testes and is investigated in relation to neurodegenerative diseases^18–20^. In different cancers, however, upregulated UCH-L1 has been linked to key cancer hallmarks including invasion^21^, metastasis^22–25^, epithelial-to-mesenchymal transition (EMT)^26^, angiogenesis^27^, cancer stemness^28,29^ and drug-mediated resistance mechanisms^27,30–32^ and correlated to poor patient prognosis^21,28,33,34^. In HCC, the role of UCH-L1 remains unclear. Despite initially reported as a potential tumour suppressor^35^, more recent omics-data show elevated UCH-L1 levels in advanced HCC^36^, renewing research interest in its possible tumour-promoting role.

Here, we evaluated the therapeutic potential of combining mTKI sorafenib (SFN) and LDN57444, the most frequently used inhibitor of UCH-L1^37–40^. *In vitro,* LDN57444 acts strongly synergistic with SFN and, at reduced SFN-dose, co-treatment completely inhibits tumour growth in an HCC mouse model relative to monotherapy. Moreover, we show that LDN57444 reverts cellular adaptive responses to SFN. Together, our findings support UCH-L1 inhibition as a promising strategy to increase SFN efficacy in aHCC with potential to mitigate adverse effects and acquired resistance.

## Materials and methods

Details on materials and methods are provided in the Supplementary data.

### Orthotopic HCC xenograft mouse model

1.5 x 10^6^ Hep3B cells in 30 µL Matrigel (Corning®)/PBS (1:1) were injected into the left-lateral liver lobe of anesthetised male NOD-SCID mice (8-12 weeks old). After twenty-four days, when tumours reached ∼100-200 mm^3^ (based on pilot studies), mice were randomised across treatment conditions and treated daily by intraperitoneal injection with vehicle, 5 mg/kg LDN57444, 15 mg/kg SFN or the combination (formulation: see Supplementary data). Mouse body weight was monitored twice weekly. After 3 weeks of treatment, mice were sacrificed, excised tumours weighed, measured using a caliper and processed for downstream analyses (Western Blot, RT-qPCR, immunohistochemistry). Tumour volume was calculated (length (mm) x width^2^ (mm) x 0.5).

### Quantitative analysis of spheroid invasion

3D-matrix-embedded Hep3B spheroids were drug treated and imaged (Supplementary data). To compare treatment effects, SImBA–SiQuAl extracted quantitative spheroid features (morphologic, growth/invasion kinetics, viability) and performed multiparametric PCA, k-means clustering and statistics. By identifying the features driving the clustering, differences in phenotype were uncovered (for details: bioRxiv 2026.04.14.718366).

## Results

### Identification of a ‘high UCH-L1’ subset of HCC patients

To gain insight in UCH-L1 expression in HCC patients, we analysed TCGA-Liver Hepatocellular Carcinoma cohort data. This shows marked *UCH-L1* upregulation (*3.40 log2 Fold Change (FC)*) in primary tumours versus non-tumour liver control samples at mRNA level (Fig. 1A). Following tumour classification based on *UCH-L1* expression, Kaplan–Meier analysis revealed significantly worse overall survival (OS) for the patient subset with high *UCH-L1* tumour expression compared to the low-expression group (*p=0.02*) (Fig. 1B). Consistently, Cox proportional hazards modelling confirmed that the *UCH-L1*-high group significantly associated with poorer OS (*Hazard Risk 1.64, 95% CI: 1.08–2.49, p=0.021*), demonstrating a 64% increased mortality risk relative to the *UCH-L1*-low group.

**Fig. 1.**
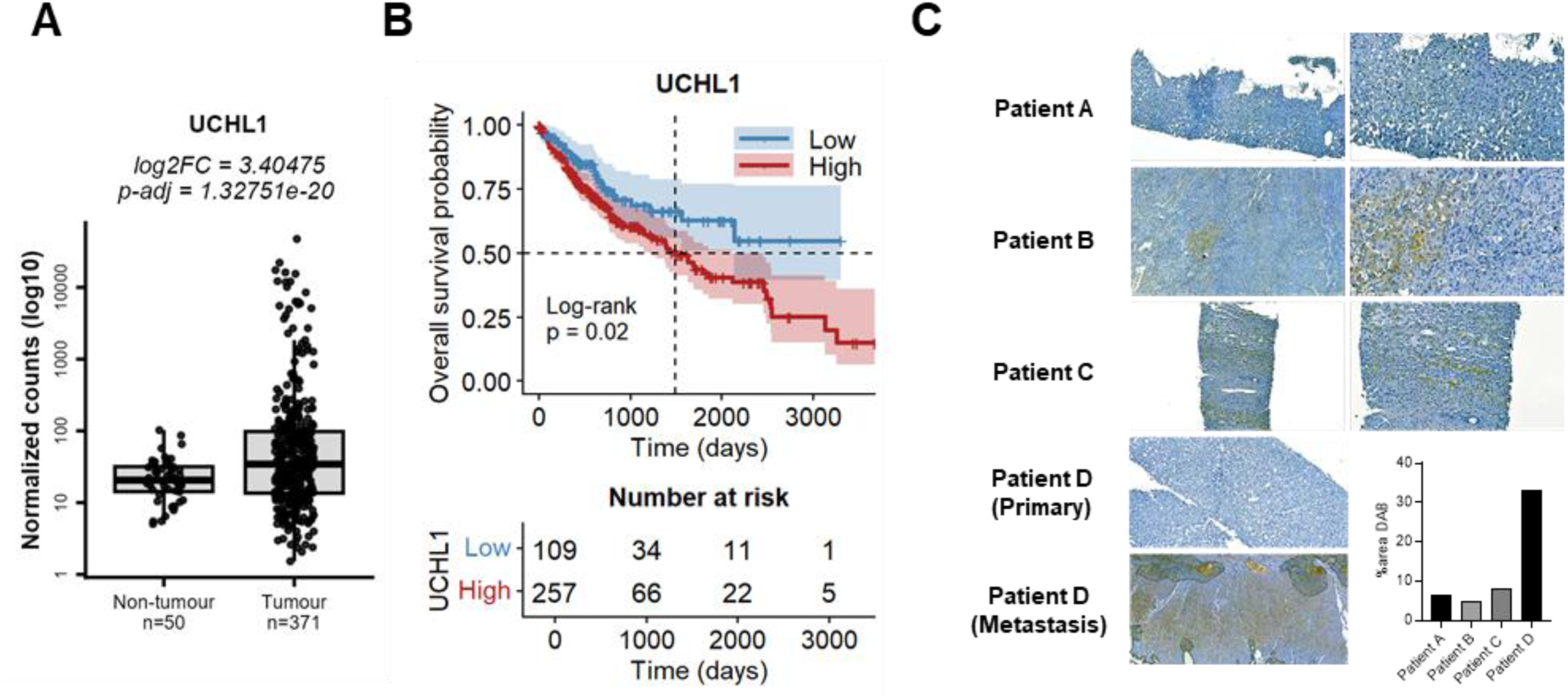
Elevated UCH-L1 expression in HCC patient tumours. (A) UCH-L1 gene expression in TCGA-LIHC cohort: log_10_(DESeq2-normalized counts) for non-tumour liver tissue and primary tumour samples. (B) Kaplan-Meier analysis of OS in TCGA-LIHC cohort categorised by UCH-L1 expression; mean OS and OS time (dashed lines); patient number at risk in time indicated. (C) UCH-L1 staining in aHCC patients tumour samples. Patient A-C: primary tumour; patient D: primary tumour and metastatic peritoneal HCC lesion (collected at diagnosis and 4 years post-diagnosis, respectively); zoom of original image indicated. Quantification of the immune response based on DAB staining is shown (for patient D: metastatic lesion).

We next performed immunohistochemical staining of UCH-L1 on patient tumour specimens. Three out of five patients showed primary tumours that were UCH-L1 positive (patient A-C), albeit with substantial inter- and intratumoral heterogeneity in immunostaining (Fig. 1C). Notably, for a patient whose primary tumour lacked UCH-L1 expression (patient D), a later peritoneal metastatic lesion contained extensive regions of UCH-L1 positive cells, with overall expression levels exceeding those observed in the UCH-L1 positive primary tumours (Fig. 1C). Together with the results from TCGA-LIHC, this indicates that UCH-L1 is upregulated in a subset of HCC patients.

### Subtoxic doses of sorafenib induce UCH-L1 expression and promote invasiveness in Hep3B cells *in vitro*

To document UCH-L1 levels in SFN-treated HCC cells, we first determined the *in vitro* cytotoxicity profile of SFN over a broad concentration range (0-35 µM, 24h) in Hep3B cells (Fig. 2A). SFN displayed a biphasic dose-response curve, characterized by a first EC_50_ of 3.6 µM (∼80% toxicity) and a clear plateau (8-14 µM, ∼60% toxicity). At low SFN doses (1 and 3.6 µM), which induced no or low cytotoxicity, UCH-L1 protein levels were significantly upregulated (∼2.5-4-fold) (Fig 2B). Higher SFN concentrations (≥ 12 µM) did not further increase UCH-L1 levels (Fig. S1A). UCH-L1 upregulation by low-dose SFN was confirmed using immunostaining (Fig. 2C).

**Fig. 2.**
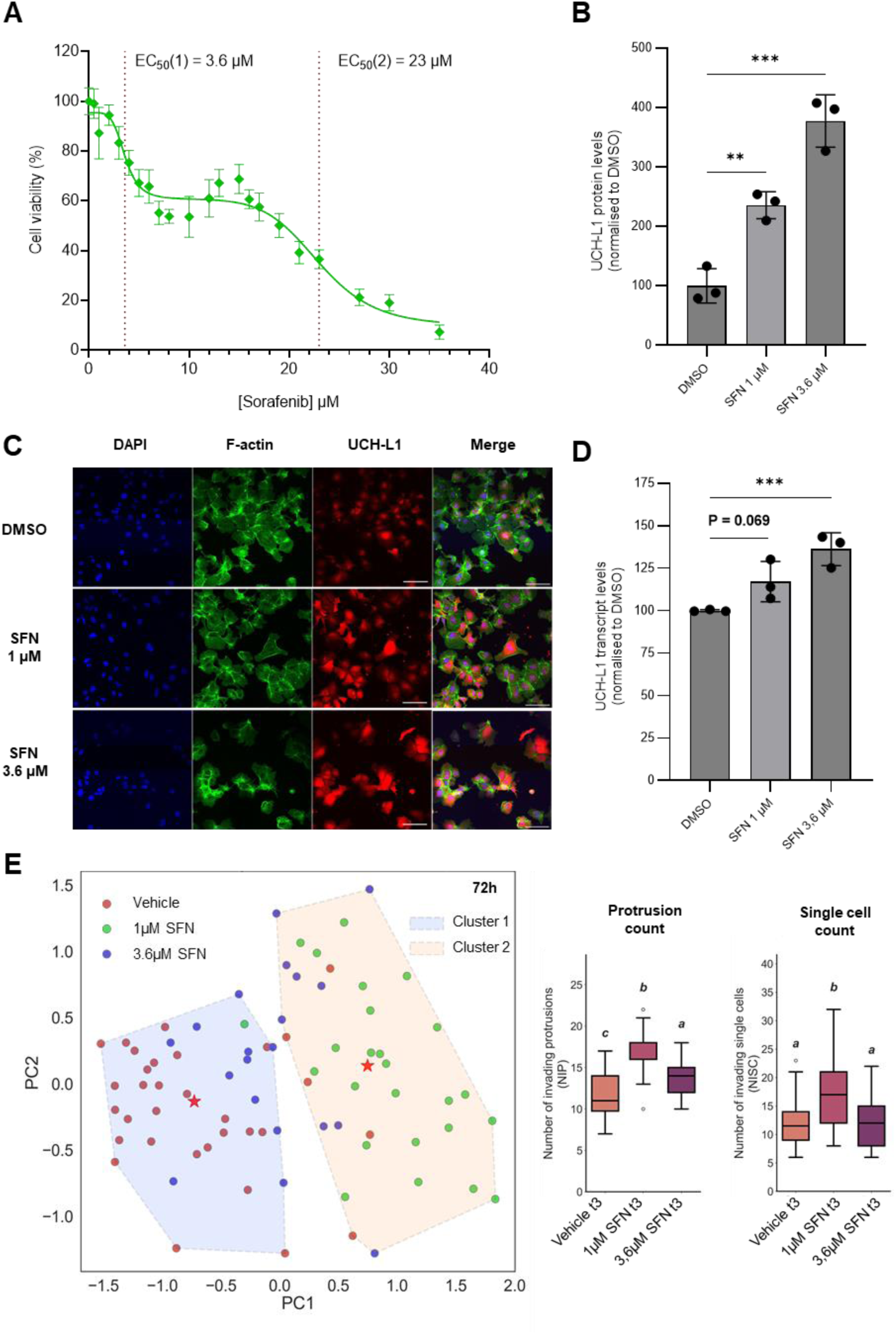
Low-dose SFN induces UCH-L1 upregulation and promotes invasiveness in Hep3B cells. (A) Biphasic dose-response of SFN on Hep3B cell viability; red-dotted lines: EC_50_-values. (B) UCH-L1 protein levels following 24h low-dose SFN, relative to control (DMSO). (C) Representative immunofluorescent images of control or 24h SFN-treated Hep3B cells; DAPI (blue), F-actin (green), UCH-L1 (red). (D) UCH-L1 mRNA levels following 24h low-dose SFN, relative to control (DMSO). (B,D) mean ± SD (n=3); \**p*<0.05, \*\**p*<0.01, \*\*\**p*<0.001, \*\*\*\**p*<0.0001 (one-way ANOVA with post-hoc testing). (E) Effect of 72h low-dose SFN on phenotype of matrix-embedded spheroids; (left) PCA + k-means-clustering with distinct phenotype for 1 µM SFN (green); dots: spheres with n=32 (DMSO), 25 (1 µM SFN), 20 (3.6 µM SFN); (right). Median and interquartile range (IQR) for two spheroid features; t3: 72h; a,b,c: significantly different groups p<0.05 (ANOVA/Tukey or Kruskal–Wallis/Dunn).

UCH-L1 mRNA was also significantly increased in Hep3B cells at low-dose SFN (Fig. 2D). Analysis of publicly available RNA-sequencing data supported this: in Hep3B cells, 2 µM SFN (24h) upregulated UCH-L1 transcripts (*1.14 log2 FC*; GEO GSE151412) (Fig. S1B) and Hep3B cells with acquired SFN-resistance following long-term treatment displayed an even greater increase in UCH-L1 transcripts compared with vehicle-treated cells (*2.49 log2 FC*; GEO GSE158458) (Fig. S1C). Notably, in Hep3B, the low, non-toxic 1 µM SFN dose also induced a significant elevation in Nrf-2 protein (Fig. S2), a reported transcriptional regulator of *UCH-L1*^41^.

Motivated by the SFN-induced UCH-L1 upregulation at low doses and the association of elevated UCH-L1 levels with aggressive disease, we next assessed whether low-dose SFN affects Hep3B invasiveness *in vitro*. Growth and invasion of 3D-matrix-embedded Hep3B spheroids treated with vehicle, 1 µM SFN or 3.6 µM SFN were imaged after 24, 48 and 72h and the drug response was determined based on an extensive spheroid feature set (SImBA-SiQuAl software, bioRxiv 2026.04.14.718366). PCA-cluster analysis showed that 1 µM SFN-treated spheroids clustered separately from control spheroids (Fig. 2E, Fig. S3A). Feature analysis indicated that this differential clustering is caused by increased invasiveness. Indeed, 1 µM SFN-treated spheroids showed more protrusions and more invading single cells (Fig. 2E), larger invasive radii, larger perimeter, reduced circularity and increased growth rates (Fig. S3B-C). Note that low-dose SFN was not cytotoxic to HCC-spheroids, since evident cytotoxicity requires 10 µM SFN or higher (Fig S3D). Interestingly, based on clustering results and individual features, spheroids treated with 3.6 µM SFN show a distinct phenotype, intermediate between control and 1 µM SFN (Fig. 2E, Fig S3A-C). This suggests that at 3.6 µM, additional SFN-dependent signalling and/or cytostatic effects are at play, partially countering the effect on invasion.

Collectively, we show that *in vitro* in Hep3B cells, SFN strongly induces UCH-L1 upregulation at non- and subtoxic doses (1-3.6 µM) and induces rapid and sustained invasion of 3D-spheroids at the non-toxic 1 µM SFN dose.

### UCH-L1 inhibition by LDN57444 strongly sensitizes HCC cells to sorafenib and impairs sorafenib-induced invasiveness

Since low-dose SFN induced UCH-L1 expression, we investigated the effect of pharmacological inhibition of UCH-L1 on SFN-cytotoxicity in 2D-cultured Hep3B cells using LDN57444 (LDN), a reversible, active site-directed small-molecule inhibitor of UCH-L1^42^. Combining non- or low-toxic LDN concentrations (Fig. S4A) with a range of SFN doses (0-35 µM) markedly increased cytotoxicity versus SFN, reducing IC_50_ for the combination to 2.1 µM (Fig. 3A). This effect was very prominent using 10 µM LDN, but already significant at 2.5 and 5 µM LDN (Fig. S4B). Synergy scoring revealed synergistic effects for low-dose combinations, with a pronounced synergy peak at 4 µM SFN + 10 µM LDN (Fig. 3B). In addition, also in 3D-Hep3B spheroids, LDN (5-15 µM) and SFN (3.6 µM) displayed strong synergistic cytotoxic effects (Fig. S4C). This highlights that LDN sensitizes to SFN and that synergy is strongest where SFN-dependent UCH-L1 upregulation is most prominent.

**Fig. 3.**
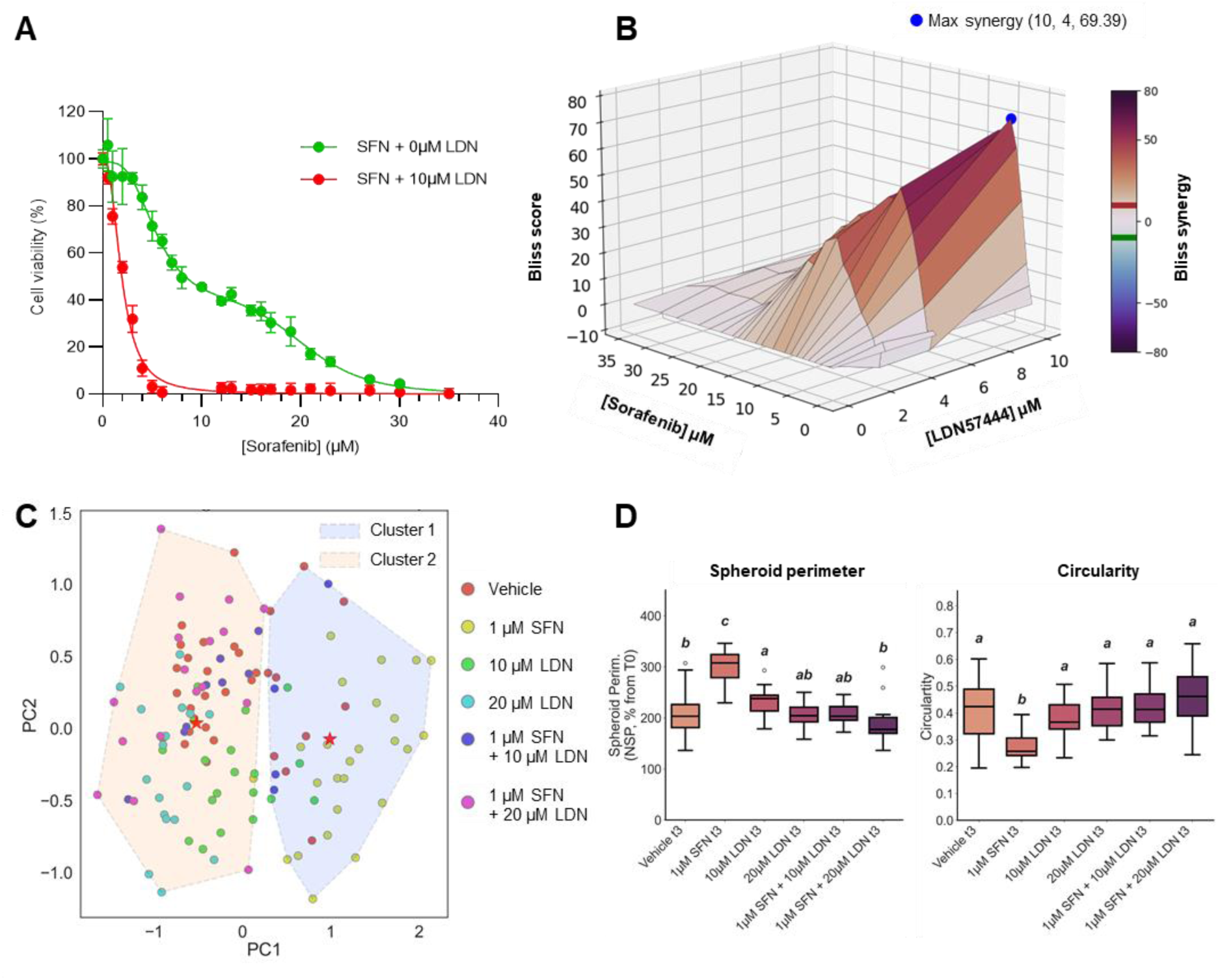
Combining UCH-L1 inhibition and low-dose SFN in Hep3B cells: synergistic cytotoxicity and suppression of SFN-induced invasion. (A) % cell viability in 2D-cultured cells after treatment with SFN alone (green, 0-35 µM) or SFN and 10 µM LDN combined (red). (B) Synergy score plot for cytotoxicity of SFN/LDN combinations. Blue dot: maximal synergy. (C) Multiparametric phenotype analysis (PCA, k-means clustering) of 72h treated 3D-spheroids. Combination-treated spheres (blue, purple dots) mainly are in cluster 2 with control spheroids (red). (D) Quantification of spheroid perimeter and circularity across treatments at 72 h (t3). Boxplots: median and IQR; a,b,c: *p*<0.05. (C,D) n/condition, 24-48h result and more features see Fig S5.

Even stronger synergistic cytotoxicity was observed in SNU-423 HCC cells. Compared to Hep3B, SNU-423 inherently have higher UCH-L1 levels that are not further induced by (low-dose) SFN (Fig. S4D-E). Strikingly, in SNU-423 cells, LDN-SFN synergy occurred at even lower SFN doses than in Hep3B: 2 µM SFN and 2.5 µM LDN, that were non-toxic on their own, produced complete cytotoxicity when combined (Fig. S4F-G). Together, these results show that UCH-L1 inhibition by LDN sensitizes HCC cells to SFN, irrespective of whether high UCH-L1 was inherent (as in SNU-423) or SFN-induced (as in Hep3B).

Next, we determined whether UCH-L1 inhibition by LDN also impacts SFN-induced invasive behaviour. Hep3B spheroids were treated with 1 µM SFN, 10 or 20 µM LDN or SFN-LDN combinations. Clearly, LDN alone had little effect: LDN treated spheroids overall behaved as vehicle controls (Fig. 3C, Fig. S5A-B, illustrated in Fig. S6). However, when LDN was combined with 1 µM SFN, this markedly affected the phenotype induced by 1 µM SFN alone. Indeed, co-treated spheroids did not cluster with 1 µM SFN but, due to the loss of the SFN-induced invasiveness, with controls and LDN-mono-treated spheroids (Fig. 3C, Fig. S5A, Fig. S6). For co-treated spheroids, features such as spheroid perimeter and circularity were indeed consistently significantly different from the 1µM SFN condition, and mostly not significantly different from the controls (Fig. 3D, Fig. S5B). Importantly, this loss of invasion upon co-treatment appeared not attributable to cytotoxicity since neither LDN nor combinations with 1 µM SFN induced substantial toxicity levels (Fig. S5C, compare to cytotoxicity level in Fig. S3D for high SFN and in Fig. S4C for combination at higher SFN). Together, these data demonstrate that LDN-mediated UCH-L1 inhibition effectively blocks the pro-invasive behaviour induced by low-dose SFN, suggesting a possible role for SFN-induced UCH-L1 upregulation as driver of this pro-invasive response.

### LDN57444 reverses sorafenib-driven MAPK activation and oncogenic signatures in HCC cells

To gain insight into the molecular basis of the low-dose SFN response and the associated LDN-mediated sensitisation and invasion suppression, we examined MAPK-pathway activity in Hep3B cells, motivated by previous reports describing paradoxical MEK1/2-ERK1/2 reactivation following *in vitro* SFN treatment^43,44^. Consistently, we show that low-dose SFN (1 and 3.6 µM) results in sustained MAPK-activation, reflected by significantly increased MEK1/2- and ERK1/2-phosphorylation and elevated MEK1/2 (not ERK1/2) protein level (Fig. 4A-B). We next examined SNAI-1 and PD-L1, proteins linked to EMT and immune tolerance, respectively and of which expression is known to rely on MEK1/2-ERK1/2-signalling^45–48^. As anticipated based on MEK/ERK-phosphorylation status, both were significantly upregulated under low-dose SFN in Hep3B cells (Fig. 4A-B). Since protein kinase C (PKC) was recently implicated in direct phosphorylation and activation of MEK1/2 in Her2-positive breast cancer^49^, we also assessed its protein levels and showed that low-dose SFN significantly increased pan-PKC levels in Hep3B cells (Fig. 4A-B).

**Fig. 4.**
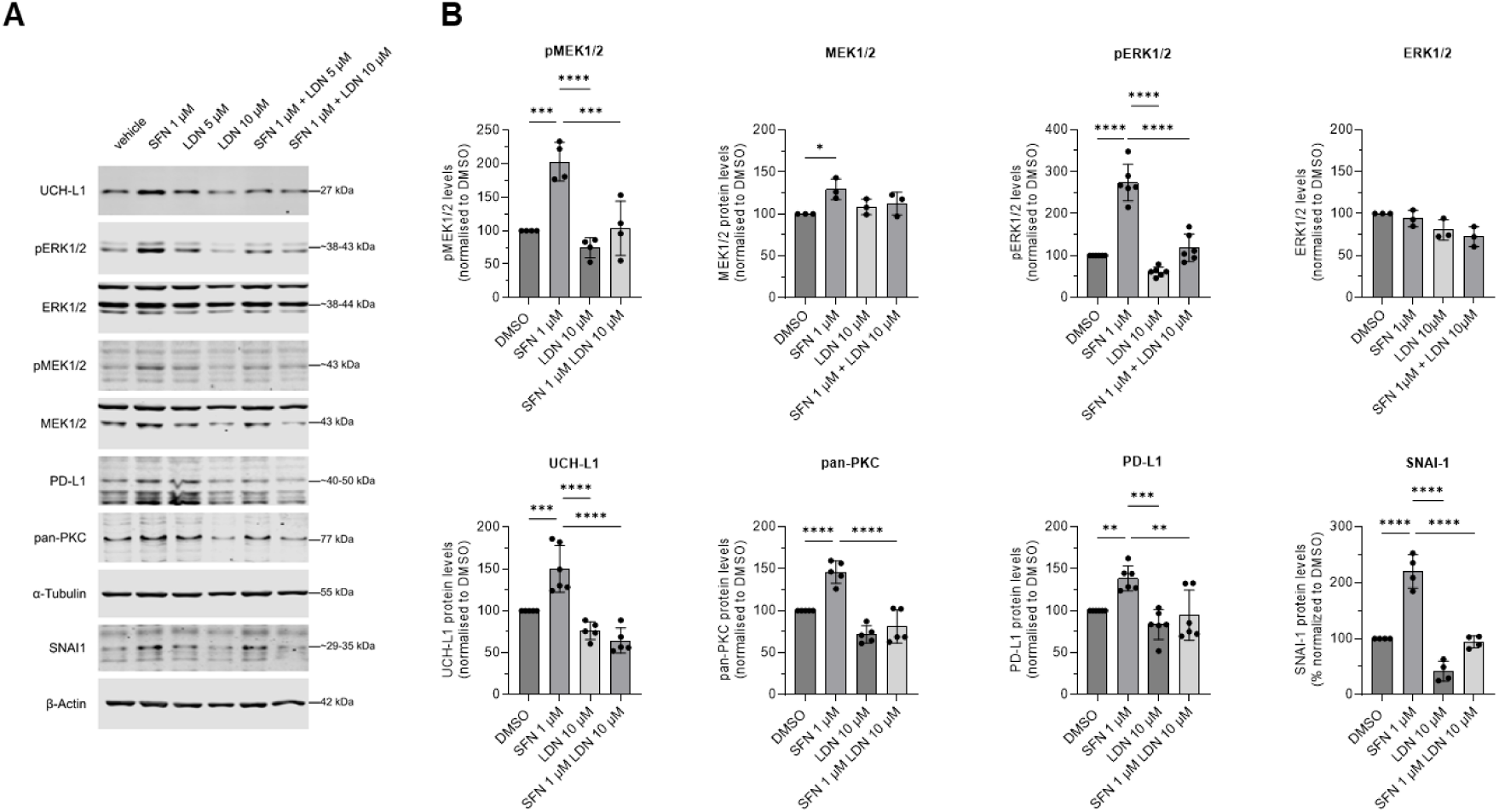
LDN-mediated UCH-L1 inhibition reverts SFN-induced sustained MAPK activation and downstream signals in Hep3B cells. (A) Representative immunoblots of Hep3B cell lysates following 24h cell treatment as indicated. α-tubulin: loading control (β-actin for SNAI-1). (B) Relative protein level in cell lysates following 24h cell treatment: vehicle (DMSO), SFN (1-3.6 µM), LDN (5-10 µM) or their combinations based on immunoblotting. Data are mean ratio over loading control ± SD (n=3-6); \**p*<0.05, \*\**p*<0.01, \*\*\**p*<0.001, \*\*\*\**p*<0.0001 (one-way ANOVA with post-hoc testing).

Next, we evaluated whether LDN treatment affects the adaptations induced by low-dose SFN in Hep3B. Strikingly, SFN co-treatment with 5 or 10 µM LDN completely reverted the MEK1/2-ERK1/2 activation induced by 1 µM SFN alone: lower p-MEK and p-ERK were observed and the upregulation of PKC, SNAI-1 and PD-L1 was suppressed (Fig. 4A-B). Interestingly, protein levels of the UCH-L1- transcription factor Nrf-2, were also downregulated by combining 1 µM SFN and LDN (Fig. S2). Together with the SFN-dependent UCH-L1 upregulation, these *in vitro* findings position UCH-L1 as a regulator of SFN-driven adaptive responses, and demonstrate that UCH-L1 inhibition by LDN can rewire the cellular adaptations in the MAPK-pathway, thereby enhancing SFN efficacy.

### LDN57444 and sorafenib co-treatment significantly suppresses tumour growth in an orthotopic HCC mouse model

To validate the *in vitro* results *in vivo* and evaluate the therapeutic potential of combining SFN with LDN, we utilized an orthotopic Hep3B xenograft mouse model (Fig. 5A). Orthotopic Hep3B injection effectively resulted in HCC tumour growth as evidenced by high levels of the tumoral marker heat-shock protein (HSP70), and the established HCC markers glypican-3 (GPC-3) and glutamine synthetase (GS) compared with non-HCC tumour controls (Fig. 6A).

**Fig. 5.**
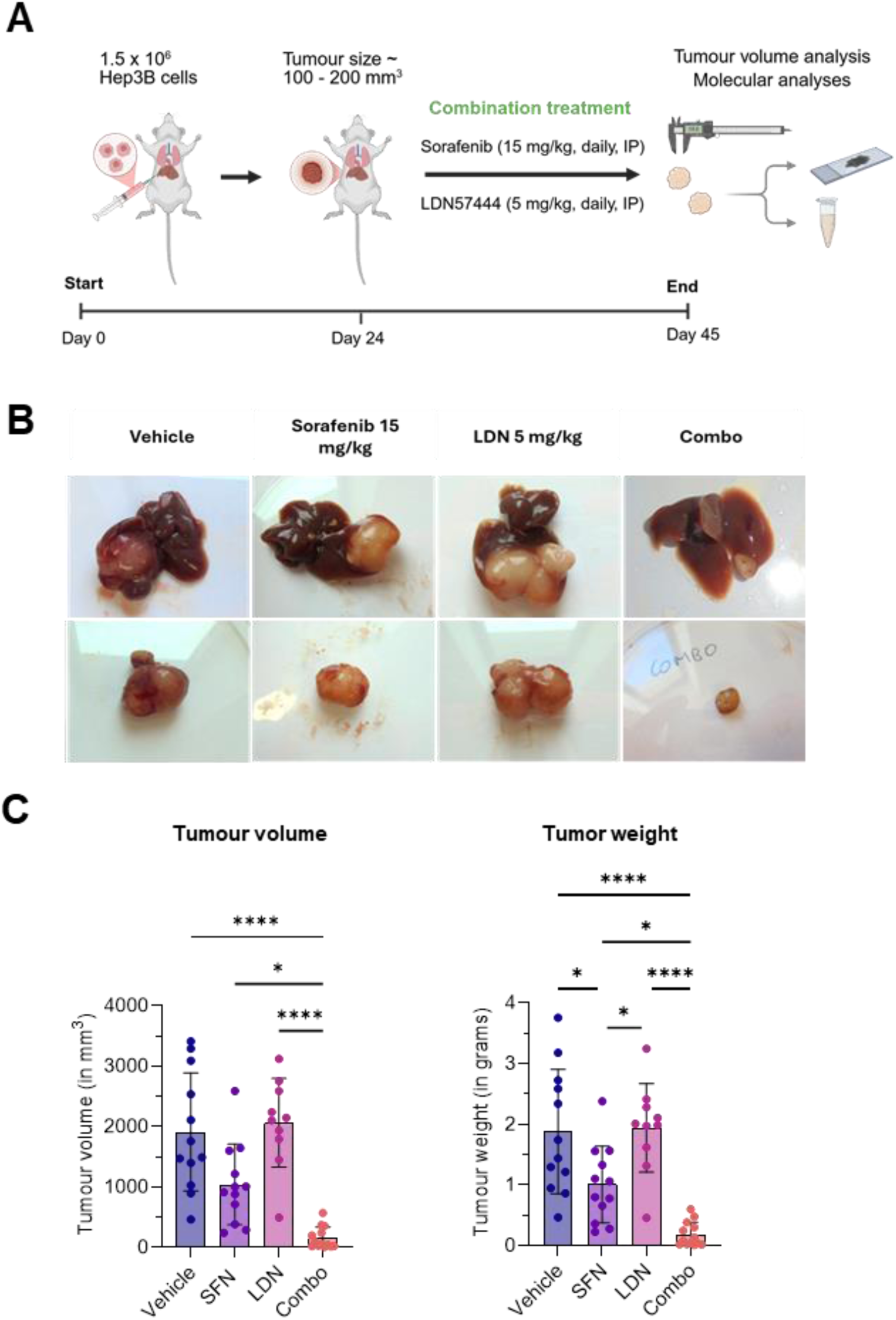
SFN-LDN co-treatment reduces HCC tumour burden in a Hep3B orthotopic xenograft mouse model at low SFN dose. (A) Workflow and treatment regimen. (B) Representative images of liver-embedded Hep3B xenograft tumours (upper) and excised tumours (lower) from treatment groups after 3-week treatment. (C) Tumour burden by volume and weight. Data are mean ± SD; vehicle (n=12), SFN (n=12), LDN (n=10), combo (n=13). \**p*<0.05, \*\**p*<0.01, \*\*\**p*<0.001, \*\*\*\**p*<0.0001 (one-way ANOVA with post-hoc testing).

**Fig. 6.**
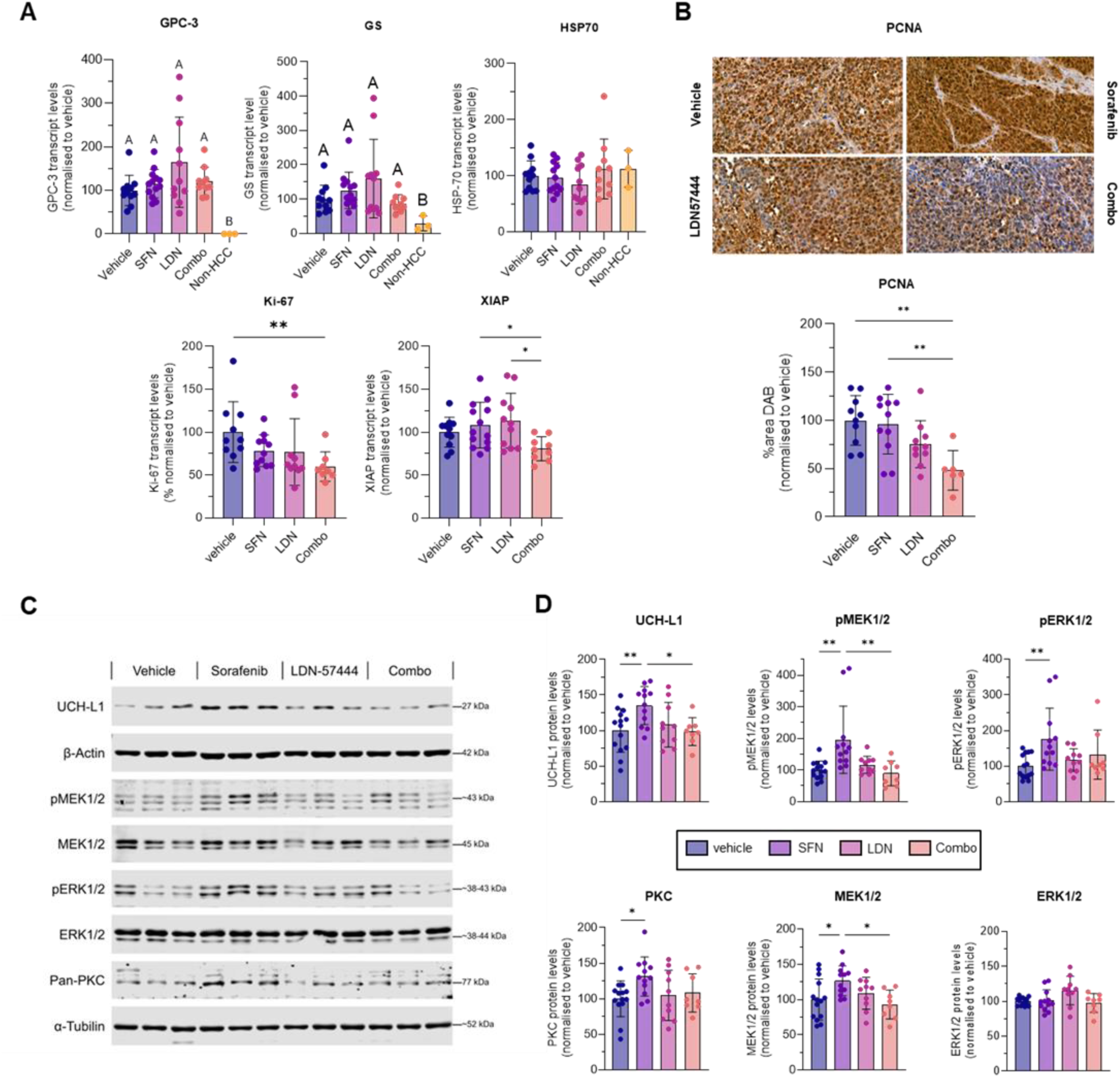
SFN-LDN co-treatment reverts SFN-induced MAPK activation and suppresses tumour proliferative and pro-survival signals in Hep3B orthotopic xenografts. (A) Relative mRNA levels of Glypican-3, Glutamine synthase, HSP70, Ki-67 and XIAP measured by RT-qPCR in homogenized Hep3B tumour tissue from mice treated with vehicle (n=10-11, set to 100%), SFN (15 mg/kg) (n=11-13), LDN (5 mg/kg) (n=10-11) or their combination (indicated as ‘combo’) (n=8-10). Data are mean ± SD; non-HCC (n=3). (B) Representative immunohistochemical staining for PCNA in Hep3B xenograft sections across treatment groups with quantification of % DAB area; vehicle (n=10, set to 100%), SFN (n=11), LDN (n=10), combo (n=6). (C) Representative immunoblots of tumour lysates from vehicle-, SFN-(15 mg/kg), LDN-(5 mg/kg) or combination-treated xenografts. β-actin: loading control for UCH-L1; α-tubulin for pan-PKC, MEK1/2, ERK1/2, phospho-MEK1/2 and –ERK1/2. Three biological replicates per condition shown. (D) Relative protein levels in Hep3B xenograft tumour lysates based on immunoblotting. Data are mean ratio over loading control ± SD; vehicle (n=7), SFN (n=6), LDN (n=5), combo (n=4); experiment performed in duplicate; (A,B,D) \**p*<0.05, \*\**p*<0.01, \*\*\**p*<0.001, \*\*\*\**p*<0.0001 (one-way ANOVA with post-hoc testing).

Tumour-bearing mice were treated daily for three weeks (Fig. 5A). In an initial pilot experiment, we compared the anti-tumour effects of combining 5 mg/kg LDN with 30 mg/kg SFN to mono-treatments. While LDN alone had no effect on tumour burden, both SFN monotherapy and the combination produced comparable near maximal tumour growth inhibition (Fig. S7A) indicating that, in this set-up, a beneficial effect of the combination versus SFN could not be assessed. Strikingly, reducing the SFN dose by half (15 mg/kg) produced markedly different outcomes. Whereas 15 mg/kg SFN only moderately inhibited HCC tumour growth, co-treatment with LDN (5 mg/kg) maximally enhanced the SFN effect indicative of a strong *in vivo* drug synergy. Tumour volumes and weights were not only significantly lower in the SFN-LDN combination compared to monotherapies (Fig. 5B-C) but, in addition, the combination achieved complete tumour growth inhibition relative to baseline tumour volume at treatment initiation (Fig. S7B-C). More so, the inhibitory effect on tumour growth obtained using LDN+15 mg/kg SFN was as strong as for 30 mg/kg SFN monotherapy (Fig. 5C, Fig. S7A). This demonstrates that high SFN efficacy can be reached at substantially lower SFN doses when combined with LDN-mediated UCH-L1 inhibition.

Co-treated tumours exhibited significantly lower Ki-67 and XIAP transcript levels compared with vehicle and monotherapies, indicating reduced proliferation and pro-survival activity (Fig. 6A). Immunohistochemistry corroborated these findings, demonstrating significantly lower proliferative cellular nuclear antigen (PCNA) levels in co-treated tumours versus SFN monotherapy and vehicle (Fig. 6B).

We next analysed by Western blot in Hep3B tumour lysates whether our *in vitro* findings on alterations in the MAPK-pathway (Fig. 4A-B) were mirrored *in vivo*. Consistently, SFN monotherapy significantly increased UCH-L1, pan-PKC, MEK1/2 (not ERK1/2) protein levels and also MEK1/2-ERK1/2 phosphorylation, indicative of sustained MAPK-pathway activation *in vivo* (Fig. 6C-D). Importantly, SFN-LDN co-treatment reverted SFN-induced upregulation of UCH-L1 protein and possibly (p)MEK/pERK (Fig. 6C-D).

Together, the *in vivo* observations align with *in vitro* synergy findings (Fig. 3A-B), and support the use of LDN-mediated UCH-L1-inhibition to disrupt SFN-driven adaptive responses in HCC.

## Discussion

aHCC remains a major clinical challenge, as current therapies provide only limited durable responses. Their clinical benefit is undermined by resistance mechanisms and dose-limiting toxicities, underscoring an unmet need for innovative strategies. Here, we provide preclinical evidence that combining SFN with LDN57444, the most used pharmacological UCH-L1 inhibitor, holds potential to address this therapeutic need.

In HCC patients, a subset of cases presents with upregulated UCH-L1 expression, associated with worse OS, supporting the existence of a clinically relevant UCH-L1-high state^36,50^ (Fig. 1). Likewise, UCH-L1 is strongly upregulated in SFN resistant HCC cells^51^ (Fig. S1). Moreover, we show that SFN induces UCH-L1 expression in UCH-L1-low HCC cells, particularly at lower doses, and that a comparable upregulation occurs in orthotopic xenografts from SFN-treated mice. Our data additionally demonstrate (i) that low doses of SFN induce an adaptive response in HCC cells characterised by increased UCH-L1 levels and (ii) that this SFN-induced adaptation is countered by LDN-based UCH-L1 inhibition resulting in enhanced SFN efficacy (as outlined in Fig. 7).

**Fig. 7.**
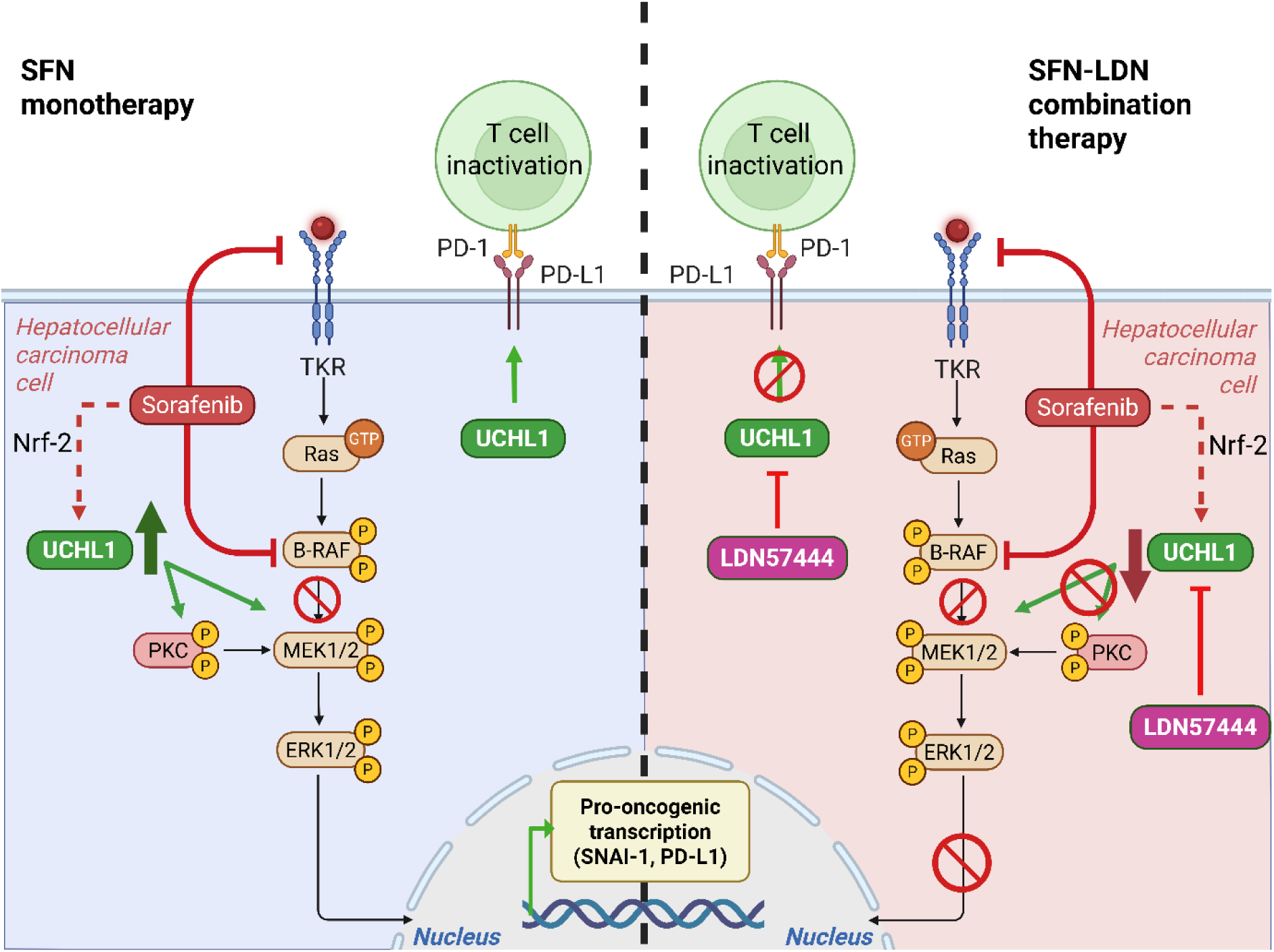
Schematic representation of the proposed role for UCH-L1 and UCH-L1-inhibition in SFN treated HCC cells: (left) Low-dose SFN induces compensatory UCH-L1 upregulation, which in turn sustains active MEK1/2-ERK1/2 signaling; (right) Upon co-treatment with SFN, adaptive responses to SFN involving UCH-L1 are re, MAPK-activation and downstream effects are inhibited, thereby enhancing SFN efficacy.

We show that SFN-induced UCH-L1 upregulation *in vitro* coincides with a broader set of drug-induced cellular responses (Fig. 7, left). First, this coincides with upregulation of Nrf-2, consistent with activation of a coordinated cellular stress adaptation as Nrf-2 is a well-established mediator of SFN-resistance and cell-survival^52^. A recent study identifying UCH-L1 as a direct transcriptional target of Nrf-2^41^, supports a possible Nrf-2-driven regulation of the SFN-induced UCH-L1 expression. Second, we demonstrate that both by low-dose SFN *in vitro* and in SFN-treated HCC xenograft tumours, UCH-L1 upregulation co-occurs with sustained activation of MEK1/2-ERK1/2-signalling and increased PD-L1 and SNAI-1 levels (Fig. 7, left). Together, these features are indicative of activated oncogenic signalling promoting proliferation, survival and invasion. Similar paradoxical MEK1/2-ERK1/2-activation in response to SFN or other RAS-RAF-MEK inhibitors is increasingly reported in cancer, including HCC^44,53,54^, where it is propagated as a prominent element in acquired resistance to SFN in patients^55^. Consistently, UCH-L1 has been shown to drive PD-L1 expression and SFN-induced PD-L1 upregulation has been linked to ERK1/2-mediated NF-κB signalling^44,56^. Sustained MEK1/2-ERK1/2 signalling may also be reinforced by increased PKC expression, which also occurs under low-dose SFN (Fig. 7, left). PKC activity has been proposed as an alternative route for MEK-activation in the context of drug-resistance in cancer^49^. Finally, our data show that the cellular reprogramming induced by low-dose SFN in Hep3B cells is coupled to increased invasion *in vitro*. SFN-induced HCC invasiveness *in vitro* has previously only been reported by Zhuang et al.^57^. Importantly, elevated UCH-L1 expression has been linked to tumour invasion and metastatic progression in multiple cancers, through promoting EMT and pro-survival signalling^21,22^.

Strikingly, we show that all of these observed SFN-induced adaptive cellular responses are countered by combining a low dose of SFN with a non-toxic dose of the UCH-L1 enzymatic inhibitor LDN57444 (Fig. 7, right). Indeed, in Hep3B cells, the SFN-LDN combination suppresses MEK1/2-ERK1/2 signalling and restores levels of PKC, SNAI-1, PD-L1, Nrf-2, and also UCH-L1 itself, to baseline and reinstates, even reinforces, the inhibitory effect of SFN. Additionally, 3D-spheroids co-treated with a low SFN dose and LDN lose their invasive potential. These findings suggest a model in which Nrf-2-mediated UCH-L1 induction forms part of an adaptive stress response to SFN, with UCH-L1 driving constitutive MEK1/2-ERK1/2 activation and promoting invasion. Notably, the currently reported UCH-L1 interactome (BIOGRID) a.o. includes MEK2, PKC2, PD-L1, and a regulator of Nrf-2, corroborating a role for UCH-L1 in these cellular adaptations.

Significantly, the effects of LDN/SFN co-treatment on cellular signalling give rise to strong SFN-sensitizing effects of the co-treatment on HCC cell viability and proliferation *in vitro* (2D, 3D) and *in vivo*. *In vitro*, we demonstrate strong synergistic cytotoxicity of the drug combination in two HCC cell lines, even at SFN-doses that are non-toxic as mono-treatment (Fig. 3, Fig.S4). *In vivo*, the drug combination enhances SFN efficacy to inhibit HCC growth at reduced SFN dose (15 mg/kg): it outperforms effects of SFN monotherapy and, when combined with LDN, this low SFN dose matches the efficacy of doubling the SFN monotherapy dose (Fig. 5, Fig. S7).

These findings may be highly clinically relevant by offering considerable therapeutic advantages. First, the tumour-restricted nature of elevated UCH-L1 expression implies that UCH-L1 targeting may exert lower systemic toxicity compared with direct inhibition of ubiquitous targets. The importance hereof is underscored by counterexamples of combining RAF- and MEK-inhibitors, which display substantial all-grade and ≥3 grade toxicities in phase-III clinical trials^58^, or for which phase-I trials were prematurely terminated due to dose-limiting adverse effects^59^. Thus, the UCH-L1 tumour-restricted expression pattern possibly offers the advantage of tumour-selective treatment effects. Secondly, the sensitizing nature of LDN to SFN *in vivo*, suggests that, upon clinical implementation, substantially lower and thus better-tolerated SFN doses could be used. As such, SFN-LDN co-administration may also lower the risk for drug-induced adverse effects. Finally, the presented co-treatment may prove valuable for future anti-resistance strategies, since the adaptive responses to SFN countered by SFN-LDN in HCC cells are known to contribute to drug resistance and relapse in patients^55^.

Our data show in both *in vitro* and *in vivo* HCC robust sensitizing effects to low SFN dose by UCHL1-inhibitor LDN. Numerous *in vitro* studies across different biological and pathological contexts support UCH-L1-dependent LDN-activity^27,37,60^. Even so, as with most small-molecule inhibitors, some reports discuss LDN target-specificity^61^, suggesting its activity may not be mediated exclusively through UCH-L1. Additional mechanistic studies will thus be needed to fully delineate the molecular framework through which LDN-based UCH-L1-targeting modulates the SFN response and acquired resistance in HCC. In addition, given the biological limitations of the preclinical models used in this study, future validation via clinical studies will be required.

In conclusion, this study proposes UCH-L1 as a regulator of adaptive oncogenic signalling triggered by SFN. Given the potency of LDN in sensitizing HCC cells to SFN, both *in vitro* and *in vivo*, our work highlights a promising opportunity to integrate small molecule-based inhibition of UCH-L1 into combination strategies for the treatment of advanced and putatively drug-resistant HCC.

## Abbreviations

DUB: deubiquitinase; ECM: extracellular matrix; HGF: hepatocyte growth factor; IP: intraperitoneal; IQR: inter-quartile range; LDN: LDN57444; OS: overall survival; SFN: sorafenib; UCH: ubiquitin C-terminal hydrolase.

## Acknowledgements

We warmly thank lab technicians Petra Van Wassenhove, Inge Van Colen, and Els Van Deynse for their excellent technical support and all members of our labs and collaborating scientists for helpful insights and discussions. We also thank the scientists/clinicians at the pathology unit of UZ Ghent for providing us with HCC patient specimens.

## Author contributions

Conception and design: MVT, HVV, LD, EVDV, KD; Supervision: MVT, LD, HVV; Animal experiments: KD, PVW, EVDV, ZDV; *In vitro* experiments and data-analysis: EVDV, KD, DB; Critical revision for important intellectual content: MVT, LD, HVV; Writing original draft: KD, EVDV; Writing review and editing: CA, MVT, LD, HVV; All authors reviewed and approved the manuscript.

## Supplementary Material and Methods

### TCGA transcriptome and survival analyses

Patient RNA-sequencing data for hepatocellular carcinoma were obtained from The Cancer Genome Atlas liver hepatocellular carcinoma cohort (TCGA-LIHC) using the TCGAbiolinks R-package. Clinical annotation data including overall survival (OS) information were derived from TCGA clinical records using GDCquery_clinic in R.

Raw gene read counts (“STAR – Counts”) were downloaded for all available samples with open-access permission. Only “primary tumour” and “solid tissue normal” samples were retained for differential expression analysis (excluding “recurrent tumours” due to limited sample size (n=3)). Differential gene expression between primary tumour (n=371) and non-tumour liver tissue (n=50) was assessed using DESeq2. Results were reported as log2 fold changes (FC) with Benjamini–Hochberg adjusted p-values to control the false discovery rate (FDR).

Survival analysis was restricted to primary tumour samples for which UCH-L1 expression and clinical outcome data were available (366/371). Note that these include HCC of all stages. Patient-normalized UCH-L1 expression values from DESeq2-normalized tumour samples were stratified into UCH-L1-high and –low expression groups using the R-function *surv_cutpoint*. This uses *maxstat* to determine the threshold that maximally separates overall survival distributions based on the log-rank statistic while enforcing a minimum group size of 25%.

OS distributions were estimated using Kaplan–Meier analysis. Median OS time and 95% confidence interval were derived where estimable (not applicable in UCH-L1-low group). Note that the OSUnivariate Cox proportional hazards model was fitted to quantify the association between UCH-L1 expression and mortality risk, reported as hazard ratio with 95% confidence intervals and corresponding p-values. Relative risk increase was calculated as (HR − 1) × 100%. Survival analysis was performed using the *survival* and *survminer* R packages.

### Analysis of GEO RNA-sequencing data on Hep3B

The publicly available RNA-sequencing datasets from the Gene Expression Omnibus (GEO) of Hep3B cells treated with 2µM sorafenib for 24h (GSE151412) and sorafenib-resistant Hep3B cells (GSE158458) were re-analysed in R using DESeq2. DESeq2-normalised counts were extracted to quantify UCH-L1 expression reported as log2 fold changes with Benjamini–Hochberg adjusted p-values to control the FDR.

### Cell lines and cultivation

The human liver cancer cell line Hep 3B2.1-7 (ATCC HB-8064) was purchased from Cytion (Eppelheim, Germany). SNU-423 (CRL-2238) liver cancer cells were obtained from ATCC. Hep3B cells were cultured in complete DMEM, i.e. Dulbecco’s modified Eagle’s medium (DMEM) supplemented with 10% foetal bovine serum (FBS) (TICO Europe) and 1% antibiotic/antimycotic (Invitrogen). SNU-423 cells were cultured in RPMI 1640 medium (Gibco), supplemented with 10% fetal bovine serum (FBS) (TICO Europe), 1% antibiotic/antimycotic (Invitrogen) and 1% sodium pyruvate (Gibco). Cells were maintained in a humified incubator at 37°C and 5% CO_2_ and their mycoplasma-negative status was regularly verified.

### Compounds and antibodies

Sorafenib tosylate (HY-10201A) and the UCH-L1 inhibitor LDN57444 (HY-18637) were purchased from MedChemExpress (Monmouth Junction, NJ, USA). Sorafenib and LDN57444 were prepared in 100% dimethylsulfoxide (DMSO) (Sigma-Aldrich, Saint Louis, MI, USA). Antibodies against UCH-L1 (13179) and PD-L1 (13684) were purchased from Cell Signaling Technology (Danvers, MA, USA). Antibodies against phospho-MEK1/2 (Ser218, Ser222, Ser226) (44-454G) and MEK1/2 (PA5-31917) were purchased from Thermo Fisher Scientific (Waltham, MA, USA). Antibodies against ERK1/2 (11257-1-AP), phospho-ERK1/2 (Thr202/Tyr204) (28733-1-AP), pan-PKC (12919-1-AP), Nrf-2 (16396-1-AP), SNAI-1 (13099-1-AP), PCNA (10205-2-AP), Ki-67 (27309-1-AP) were purchased from Proteintech (Rosemont, IL, USA). Antibodies against β-actin (A2228) and α-tubulin (T5168) were purchased from Sigma-Aldrich (Saint Louis, MI, USA). All compounds and antibodies are included in the CTAT-table.

### Cell viability and dose response assays

To assess cell viability based on metabolically active cells and generate treatment dose-response curves, the XTT (2,3-bis-(2-methoxy-4-nitro-5-sulfophenyl)-2H-tetrazolium-5-carboxanilide) cell proliferation kit II (Roche, 11465015001) was used. HCC cells were seeded at 1 x 10^4^ cells/200 µL of complete cell medium per well in 96-well plates. After 24 hours, cells were treated with a concentration range of either sorafenib, LDN57444 or their combination in complete DMEM. Each concentration was tested in quadruplicate (n = 4). Controls included blank wells (medium without cells) and vehicle controls containing cells with DMSO, at levels of ≤ 0.15 v/v%, matched across all conditions. After 24h treatment, the medium was replaced with 50 µL XTT reagent mixture in 100 µl complete DMEM (following the manufacturer’s instructions) and cells were incubated for 3 hours to allow for formazan formation. Absorbance was measured at 450 nm (with 595/655 nm as reference measurement) using a microplate reader. Data were analysed in GraphPad Prism 10 and viability values were expressed as percent cell viability normalized to DMSO vehicle control (set to 100%). Dose-response curves were generated from the normalized data using non-linear (biphasic) regression (X = concentration).

### Western Blot

Cells, tumour and liver tissues were lysed with a lysis buffer containing 7M urea, 2M thio-urea, 1 % v/v Triton, 32.4 mM dithiothreitol, an in-house prepared protease inhibitor cocktail and phosphatase inhibitors (1 mM Na_3_VO_4_, 10 mM NaF). Protein concentrations of cleared cell lysates were determined using the Bradford assay (Bio-Rad). Proteins were analysed by SDS-PAGE (8 or 12% gels) and transferred to nitrocellulose membranes (Amersham™ Protran®, Sigma-Aldrich (Saint Louis, MI, USA)). Membranes were blocked for 1h at room temperature with continuous shaking in a 1:1 mix of Intercept® TBS-blocking buffer (LI-COR Biosciences) and TBS. Blots were then incubated overnight at 4°C with primary antibodies diluted in the same 1:1 blocking buffer/TBS mix supplemented with Tween-20 (0,1% v/v). Antibody dilutions were as follows for: UCH-L1 (1:1000), MEK1/2 (1:1000), phospho-MEK1/2 (1:1000), ERK1/2 (1:4000), phospho-ERK1/2 (1:2000), pan-PKC (1:6000), PD-L1 (1/1000), SNAI-1 (1/600), Nrf-2 (1:6000), β-actin (1:3000) and α-tubulin (1:6000). Subsequently, membranes were washed in TBS-Tween-20 (0,1% v/v) and incubated for 45 minutes at room temperature with IRDye® anti-rabbit (800 nm) and anti-mouse (680 nm) fluorescently labelled secondary antibodies (LI-COR Biosciences). Membranes were imaged using a LI-COR Odyssey scanner and fluorescent protein band intensities were quantified using the Image Studio Lite ^TM^ software. Protein band intensities were normalized to loading controls: β-actin or α-tubulin.

The number of replicates (n) reported for the *in vitro* experiments refer to technical replicates; each independently processed on SDS-PAGE. For the *in vivo* experiments, n represents the number of biological replicates, with each sample analysed in duplicate (technical replicates; n = 2). Given the large number of xenograft tumours (n = 22), protein levels were quantified and normalised across blots (Fig. 6D) using an internal control sample. For representative images shown on a single blot (Fig. 6C), biological replicates (n = 3) per condition were randomly selected.

### Immunofluorescent staining

Hep3B cells were seeded on Type I collagen (Corning, AZ, USA) precoated glass coverslips (class 1, VWR, Leuven, Belgium), placed in 6-well plates. After 24h, cells (optimally at ∼70% confluency) were treated with vehicle (DMSO) or the indicated concentrations of SFN for 24h. Cells were then fixed for 30 min in 4% paraformaldehyde (Sigma-Aldrich) in PBS (Invitrogen). The subsequent steps included permeabilization (0,1 % Triton X-100 in 4% paraformaldehyde (PFA, Sigma-Aldrich) in PBS for 5 min, 45 min blocking in 10 w/v% bovine serum albumin (BSA (Sigma-Aldrich)) in PBS, overnight incubation at 4°C with rabbit UCH-L1 antibody (1/300 in 1 w/v% BSA in PBS), appropriate washing and 2h incubation at room temperature with a mix of goat Alexa594-coupled anti-rabbit secondary antibody and F-actin staining Alexa-488-phalloidin (ThermoFisher Scientific, both used at a 1/200 dilution in 1 w/v% BSA in PBS) and, after washing, a final brief incubation with DAPI for nuclear staining (Sigma-Aldrich). Mounted coverslips were imaged on a FluoView1000 Olympus confocal microscope using an Olympus 20x objective with additional zoom. Presented images (pixel size_xy_: 239 nm) were acquired under identical settings e.g. 3% intensity for the 559 lasera d High Voltage at 750 for the UCH-L1 channel, to allow comparison of UCH-L1 signal intensities across conditions. Both image processing and montage assembly was performed in Fiji (https://fiji.sc).

### Spheroid invasion assay

HCC spheroids of Hep3B cells were generated using the hanging drop method. Briefly, Hep3B cells were collected by trypsination and resuspended at 100.000 cells/mL in complete medium (cell suspension, CS), mixed with Methylcellulose 4000 (MC) (final concentration: 12 mg/ml) in a 4:1 CS-MC ratio, yielding a final cell concentration of 80.000 cells/ml. Drops of 25 µl (≈2000 cells) were dispensed onto the hydrophobic lid of a square plastic cell culture disc (Nunc, 240835) using a 12-channel multichannel pipette. To prevent evaporation during incubation, 10 ml PBS (Invitrogen) was added to the bottom dish before gently inverting the lid in a single motion. Hanging drops were incubated (37°C, 5% CO_2_) for 24h, after which compact spheroids were visible microscopically. Spheroids were collected and embedded in a 3D collagen type I matrix (CM) prepared in 12-well plates according to the method described in Van Troys et al.^1^ and Van De Vijver et al. (doi: 10.64898/2026.04.14.718366). The collagen hydrogel was prepared on ice by sequentially mixing Hank’s Balanced Salt Solution (HBSS, Gibco), 10x MEM (Gibco), NaHCO_3_, complete medium, Type I collagen (Corning; final concentration 1 mg/mL) and NaOH (for details see Table 1B in Van Troys et al.^1^). Each well contained a two-layer CM: a pre-polymerised (15 min, 37°C) bottom layer (350 µL), ensuring that spheroids embedded in the upper layer are not positioned near the well surface. A total of 15 spheroids of uniform shape, size and integrity were manually picked under a binocular microscope (LEICA MZ7.5) using a P1000 pipet tip, washed in medium in a 2 mL Eppendorf tube to remove residual MC and resuspended in 360 µl of cold collagen mix, which was subsequently pipetted dropwise onto the pre-polymerised bottom layer (maintained on a 37°C heat plate), ensuring adequate spatial separation of spheroids. After polymerisation of the top layer (20 min on 37°C heat plate, 40 minutes in incubator at 37°C, 5% CO_2_), 750 µl of culture medium was added to each well, supplemented and the specified concentrations of sorafenib, LDN57444 or their combination.

T_0_-images (0 h) were acquired at this stage, after which 50 ng/ml hepatocyte growth factor (HGF; Sigma-Aldrich) was added to each well to induce invasion. Spheroid growth and invasion into the 3D collagen matrix were monitored at different time points (0-72h) within a 24h-interval using phase-contrast imaging on an Olympus IX 81 microscope with Olympus CellSens imaging software, using a 10x objective, in accordance with the SImBA-SiQuAl imaging protocol described in Van De Vijver et al. (doi: 10.64898/2026.04.14.718366). After 72h, SYTOX Green (Invitrogen, marker of dead cells) was added to a concentration of 200 nM for 30 minutes to assess treatment-induced cytotoxicity.

### RNA extraction and reverse transcription quantitative PCR (RT-qPCR)

Total RNA was extracted from cells or from homogenized tumour and liver tissue using the Promega Maxwell® RSC instrument in combination with the Maxwell® RSC simplyRNA Tissue Kit (cat n°AS1340), according to the manufacturer’s instructions. RNA purity was evaluated by spectrophotometry (NanoDrop 2000) by calculating the A260/A280 and A260/A230 ratios, and by automated electrophoretic profiling using RNA ScreenTape on the TapeStation system (Agilent Technologies).

For cDNA synthesis, 1 µg RNA was reverse-transcribed using the iScript cDNA synthesis kit (Bio-Rad, cat n°1708891). Quantitative PCR was conducted by adding cDNA (25 ng per reaction) to a 384-well plate containing gene-specific primers (see CTAT-table, 0,4 µM) and SYBR Green qPCR Master mix (no-ROX; MedChemExpress), in a final volume of 8 µL. Reactions were run on a CFX Opus 384 (Bio-Rad) and data were analysed using CFX Maestro software and qBase+ (Biogazelle). ACTB, HMBS, YWHAZ and HPRT1 served as reference (housekeeping) genes (primer sequences listed in the CTAT-table). All samples were run in technical triplicates. For each analysis, all samples in which the same target or reference gene is analysed, were included in a single run, according to the principle of sample maximalization^2^, alongside a no-template control.

Cq-values were calculated using the single-threshold method within CFX Maestro. Technical replicates with a Cq-value deviating by >0.5 cycles from the other raw Cq-values for a specific sample are automatically flagged in qBase+ (Biogazelle) and were therefore excluded from further analysis. Mean Cq-values for each sample were normalised to the geometric Cq-mean of stable reference genes, as determined by GeNorm analysis in qBase+ (Biogazelle). Normalised expression values were subsequently processed and visualised in GraphPad Prism 10. Number of replicates (n) vary per condition for different genes depending on the number of samples excluded based on replicate exclusion (see above) or based on insufficient tumour material in case of relatively small tumours. Non-HCC control cDNA (Fig. 6A) is prepared from a mix of non-HCC cells of different cancer types (cell lines HeLa, MDA-MB-231 and A549).

### Immunohistochemistry

Tumour and liver tissue sections were deparaffinized in xylene and gradually rehydrated through an ethanol gradient. Antigen retrieval was performed in 1x citrate buffer at 95°C for 10-20 minutes, after which sections were cooled at room temperature for 30 minutes and then rinsed in deionized water (3 x 5 min.). Endogenous peroxidase activity was quenched using 3 v/v% hydrogen peroxide (Sigma Aldrich) for 10 min. Following additional washing steps in deionized water and in TBS supplemented with 0,05 v/v% Tween-20 (TBS-T), sections were blocked for 1 h at room temperature in 1 w/v% BSA – 5 v/v% goat serum in TBS-T. Sections were incubated overnight at 4°C with primary antibodies against UCH-L1 (1:30) or PCNA (1:500). After washing in TBS-T, sections were incubated for 30 min with HRP-conjugated anti-rabbit secondary antibody (SignalStain^®^ Boost IHC Detection Reagent, Cell Signaling). Signal detection was performed using 3,3’-Diaminobenzidine (DAB) chromogen, followed by haematoxylin counterstaining. Quantification of DAB staining was performed using Fiji (https://fiji.sc) and expressed as % DAB-positive area. For UCH-L1, DAB signal in the entire tissue section was quantified; PCNA-positive nuclei were quantified at 40x magnification.

For staining mouse tumour tissue, the number of replicates (n) varies per condition as some biological samples were excluded for a specific protein staining based on the presence of artefacts complicating DAB quantification. For HCC patient tissues, UCHL1 was analysed and quantified in samples from 5 different patients. These patients were selected based on disease stage and, where available, longitudinal samples of a same patient were included.

### In vivo drug administration and tissue sampling

To evaluate the effects of sorafenib, LDN57444 and their combination on the growth of Hep3B xenograft tumours in NOD-SCID mice (for generation of these tumours, see Materials and Methods and Fig. 5A), Drugs were first dissolved in 100% DMSO to prepare concentrated stock solutions: 25 mg/mL LDN57444, 75 mg/mL Sorafenib or the combined stock. Stocks were sequentially diluted with PEG300 (30 % (MedChemExpress, HY-Y0873), Tween-80 (5%) and distilled H_2_O (65%). Tumour-bearing mice received intraperitoneal injections of sorafenib (15 mg/kg/day, or as specified), LDN57444 (5 mg/kg/day) or the corresponding combination.

After 3 weeks of treatment, mice were anaesthetized by intraperitoneal injection of ketamine (100 mg/kg) and xylazine (10 mg/kg) and euthanized by cervical dislocation. Liver and spleen were excised and weighed. Designated liver lobes were processed for gene expression analysis (stabilized in RNA-later), protein expression analysis (Western Blot) and histology (PFA-fixed).Tumour growth inhibitory rate (%) was calculated as (1-Y/X) × 100 with X, mean tumour volume/weight at treatment initiation and Y, tumour volume/weight of treatment group.

### Animal experiments

All animals were housed in individually ventilated cages (IVC) in a temperature-controlled, specific-pathogen-free (SPF) animal facility at Ghent University Hospital. All *in vivo* procedures were approved by the Animal Ethics Committee of the Faculty of Medicine and Health Sciences, Ghent University (approvals 23/39 and 23/39aanv).

### Statistical analysis

Unless stated otherwise, statistical analyses were performed using GraphPad Prism 10 (GraphPad Software Inc., USA). Data are presented as mean ± SD. Normality of the data distribution was assessed prior to statistical testing. For datasets passing the assumptions of normality, differences between multiple groups were assessed using one-way analysis of variance (ANOVA) with subsequent Tukey’s post-hoc testing. For non-parametric datasets, group differences were evaluated using the Kruskal-Wallis test with Dunn’s post-hoc testing. Two-sided P-values <0.05 (*), <0.01 (**), <0.001 (***) and <0.0001 (****) were considered statistically significant.

## Supplementary figures

**Fig. S1.**
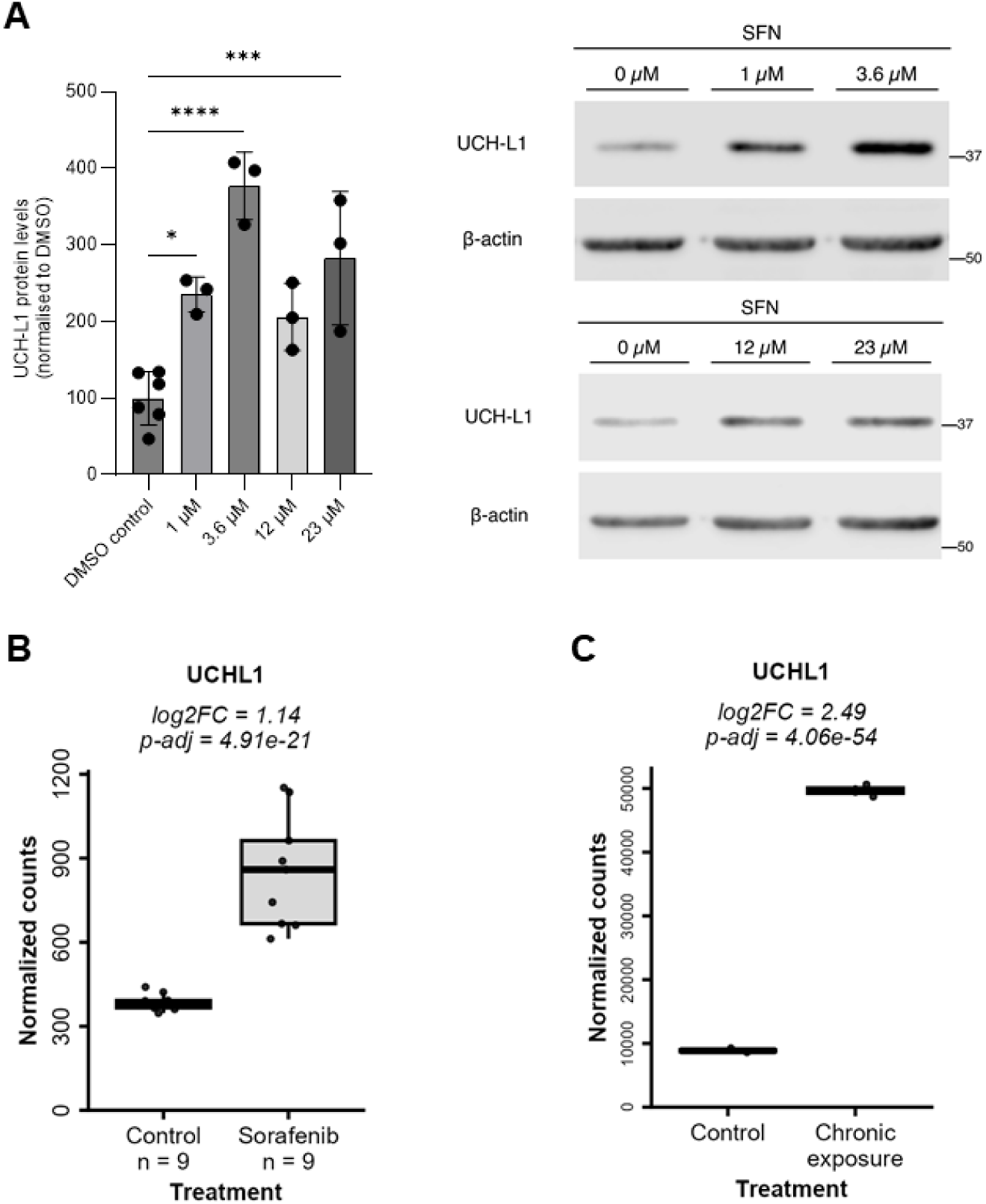
SFN induces UCH-L1 protein and gene upregulation in Hep3B cells upon short- and long-term SFN exposure. (A) Relative UCH-L1 protein levels in SFN-treated Hep3B cells (24h) normalised to DMSO control (left) with representative Western blots with β-actin as loading control (right). Hep3B cells have low basal expression (see Fig. S4E) and SFN induces UCH-L1 upregulation in these cells most prominently at low doses (1 – 3.6 µM). Bar plot data are mean ± SD; *p<0.05, **p<0.01, ***p<0.001, ****p<0.0001 (one-way ANOVA with post-hoc testing) (n=3 technical replicates per blot). (B) Upregulated UCH-L1 gene expression in Hep3B cells treated with 2 µM SFN for 24h compared to DMSO control. (GEO data GSE151412, n = 9). (C) Upregulated UCH-L1 gene expression in Hep3B cells treated with 5 µM SFN for 8 weeks compared to DMSO control (GEO data GSE158458, n = 2). (B, C) Box plots indicate median + IQR of DESeq2-normalized counts; log2 Fold Change and DESeq2 FDR adjusted p-value (Benjamini-Hochberg) are shown.

**Fig. S2.**
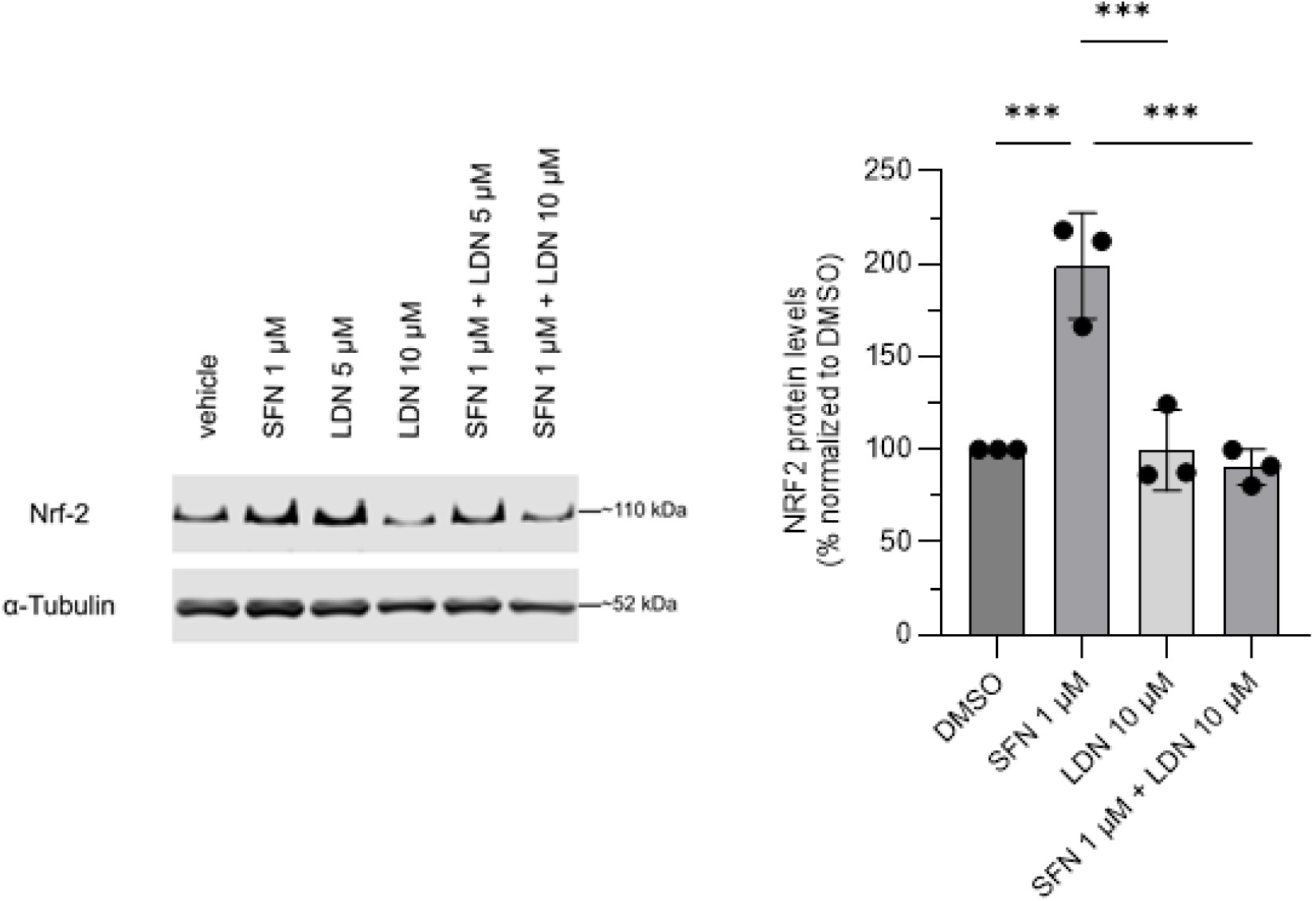
SFN-induced Nrf-2 upregulation is reverted in Hep3B cells upon co-treatment with LDN57444. Representative western blot of Nrf-2 (left) in Hep3B cells following 24h treatment with DMSO (vehicle), SFN (1 µM), LDN (5 or 10 µM), or their combinations with α-tubulin as loading control. Relative protein quantification of Nrf-2 (right) in Hep3B treated as indicated. 1 µM SFN significantly upregulated Nrf-2 protein relative to DMSO, whereas cotreatment with LDN reverted SFN-induced Nrf-2 to DMSO levels. Values are normalised to DMSO and shown as mean ± SD (n=3 technical replicates); Bar plot data are mean ± SD; *p<0.05, **p <0.01, ***p<0.001, ****p<0.0001 based on one-way ANOVA with post-hoc testing.

**Fig. S3.**
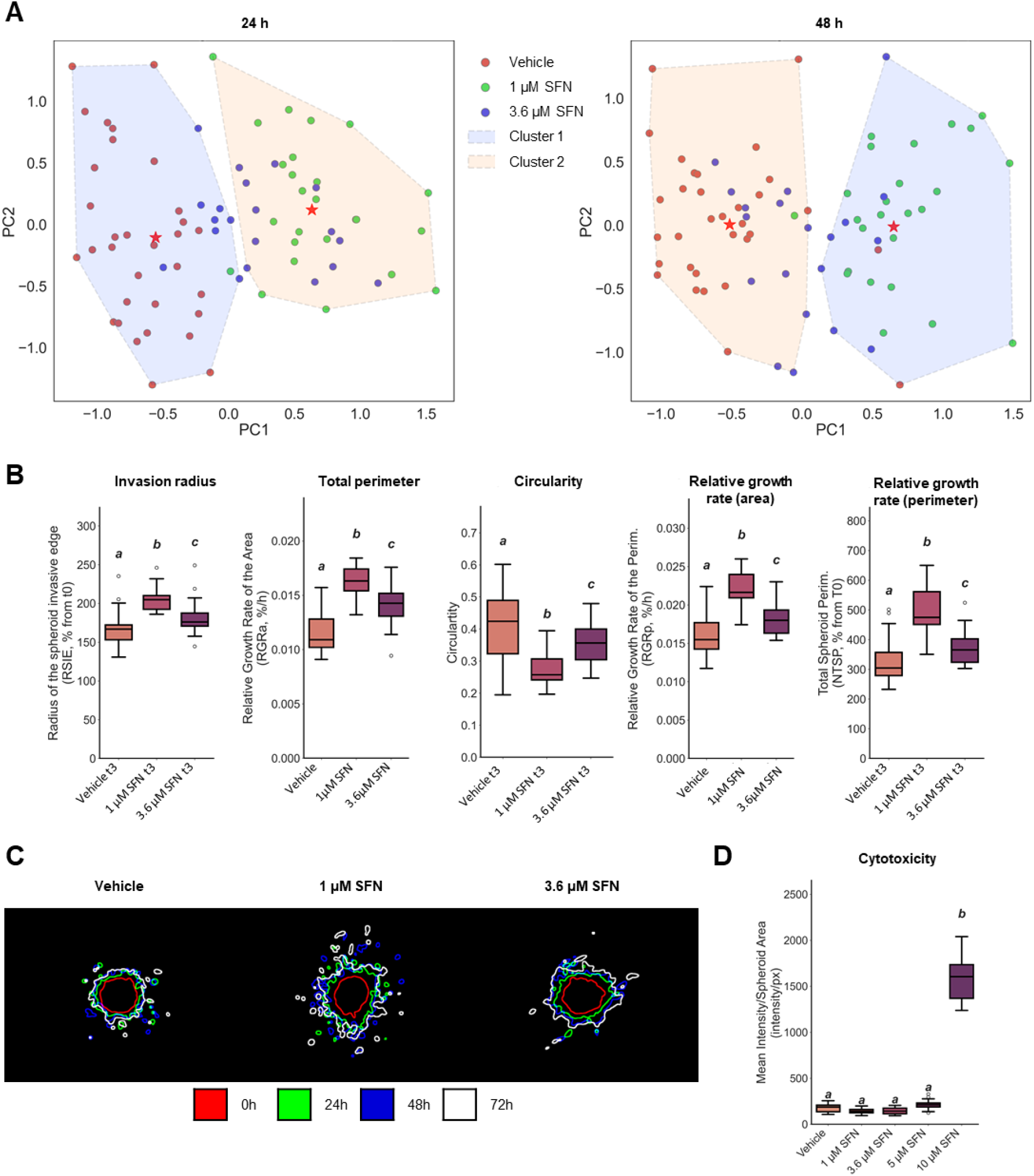
Low-dose SFN (1 µM) induces a distinct invasive spheroid phenotype, clustering separately from controls. (A) Multiparametric clustering (PCA + k-means) after 24 h (left) and 48h (right) demonstrates separate clustering for 1µM SFN already after 24h and sustained at 48h; individual dots represent spheroids, n/condition: DMSO: 32 (red), 1µM SFN: 25 (green), 3.6 µM SFN: 20 (blue). PCA-analysis is based on all SImBA-extracted growth, invasion, morphology and spheroid complexity features. (B) Boxplots (median and interquartile range) for relevant invasion and growth features after 72 h SFN treatment. The data presented is a relevant selection of all features extracted, selected to illustrate the pro-invasive phenotype induced by 1µM SFN. Included are: invasion radius (the radius of the outer bounding circle i.e. the smallest circle encompassing the spheroid and all invading cells), the total perimeter (i.e. the summed perimeter of the central spheroid mass and all invading cells/cell groups), circularity, the relative growth rate (RGR) of the area or perimeter (RGR = (ln(S_n_) – ln(S_0_))/(t_n_ – t_0_) with S_n_ the size of the spheroid area/perimeter at timepoint n (t_n_) and S_0_ the size at t_0_. (C) Overlay of spheroid outlines in time (see colours and legend) generated by SImBA of a representative spheroid per condition. Spheroids treated as indicated. 1 µM SFN-treated spheroids: more invasive phenotype. (D) Cytotoxicity in spheroids for indicated conditions at 72 h, measured as the mean intensity of SYTOX green staining within the spheroid boundary at the final imaging timepoint. No significant increase in cytotoxicity for ≤ 5 µM SFN. (B, D) Statistical significance shown as compact letter display: conditions sharing the same letter are not significantly different (p>0.05), conditions without shared letters differ significantly (p<0.05) (ANOVA/Tukey or Kruskal–Wallis/Dunn depending on normality).

**Fig. S4.**
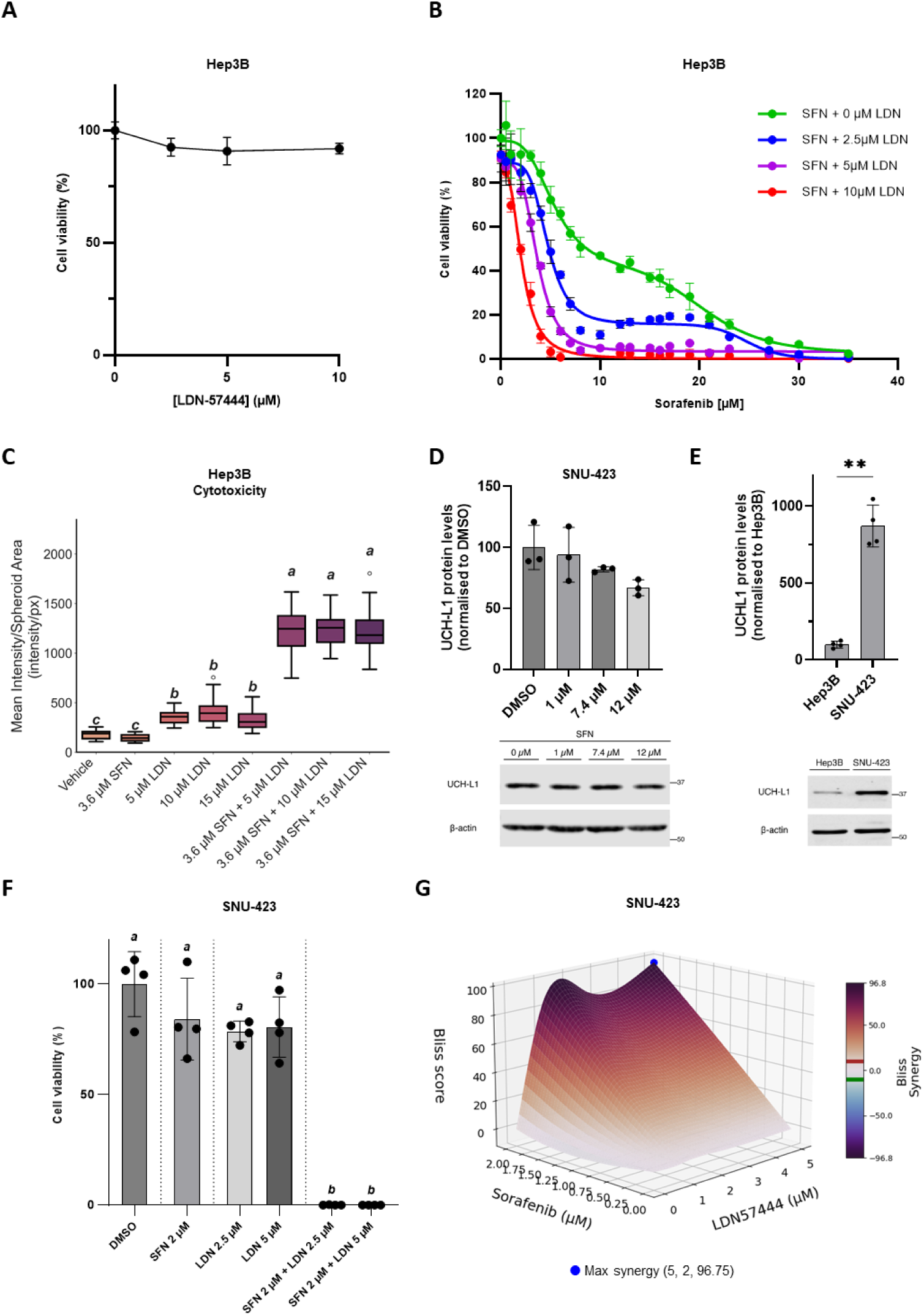
UCH-L1 inhibition by LDN sensitises Hep3B and SNU-423 cells to sorafenib cytotoxicity in a synergistic manner. (A) Cell viability of Hep3B cells cultured in 2D and treated with UCH-L1 inhibitor LDN (0-10 µM) for 24h showing minimal toxicity of LDN monotreatment across the tested range. Data based on XTT viability assay; mean ± SD (n = 4, technical replicates). (B) Dose-response curves of the SFN effect on cell viability of Hep3B cells cultured in 2D and treated with a SFN concentration range (0-35 µM) in combination with the indicated LDN concentrations (0-10µM). LDN potently sensitizes Hep3B cells to the cytotoxic activity of SFN in a dose-dependent manner, markedly reducing cell viability compared to monotreatment SFN (green curve), indicative of SFN-LDN synergy. Data based on XTT viability assay; mean ± SD (n = 4). (C) SFN-LDN synergistic cytotoxic effect on 3D Hep3B spheroids embedded in a 3D matrix displayed using boxplots (median and interquartile range). Spheroids were treated with vehicle (DMSO), 3.6 µM SFN, LDN (5, 10, 15 µM) or a combination. The number of spheroids per varies between 17 and 26. Cytotoxicity of treatment was scored after 72 h based on mean pixel intensity of SYTOX green toxicity staining in the total spheroid area. LDN induces only a slightly higher cytotoxicity than 3.6 µM SFN and vehicle in monotreatment. Hep3B spheroids treated with the SFN/LDN combinations show markedly increased toxicity, significantly higher than any other treatment and indicative of drug synergy. Statistical significance shown as compact letter display: conditions sharing the same letter are not significantly different (p>0.05), conditions without shared letters differ significantly (p<0.05) (ANOVA/Tukey or Kruskal–Wallis/Dunn depending on normality). (D) Relative UCH-L1 protein levels in SNU423 HCC cells that are 24h SFN-treated. The SFN doses used are selected based on dose response analysis (0-35 µM SFN, not shown) with 1 µM a non-toxic SFN dose. In SNU-423 cells, with high basal expression, SFN does not further induce UCH-L1 protein level (D). Bar plot data are mean ± SD; *p<0.05, **p<0.01, ***p<0.001, ****p<0.0001 (one-way ANOVA with post-hoc testing). (E) Relative UCH-L1 protein levels in untreated SNU423 cells compared to Hep3B cells. Basal expression in SNU-423 cells is 8.7-fold higher than Hep3B cells. UCH-L1 expression is normalised to Hep3B cells (n = 4; technical replicates). (D,E) A representative Western blot is shown with β-actin as loading control below the graphs. (F) % Cell viability of SNU-423 cells treated with monotreatment of SFN or LDN or with their combination at the indicated doses. In contrast to mono-treatments, SFN-LDN co-treatment results in complete loss of cell viability. This validates SFN-LDN synergistic cytotoxicity in an additional HCC cell line. (G) Interpolated Bliss synergy score surface plot identifying broad synergy of toxicity of combined treatments (LDN + SFN) in SNU-423 cells. Maximal synergy at 2 µM SFN combined with 5 µM LDN (blue dot).

**Fig. S5.**
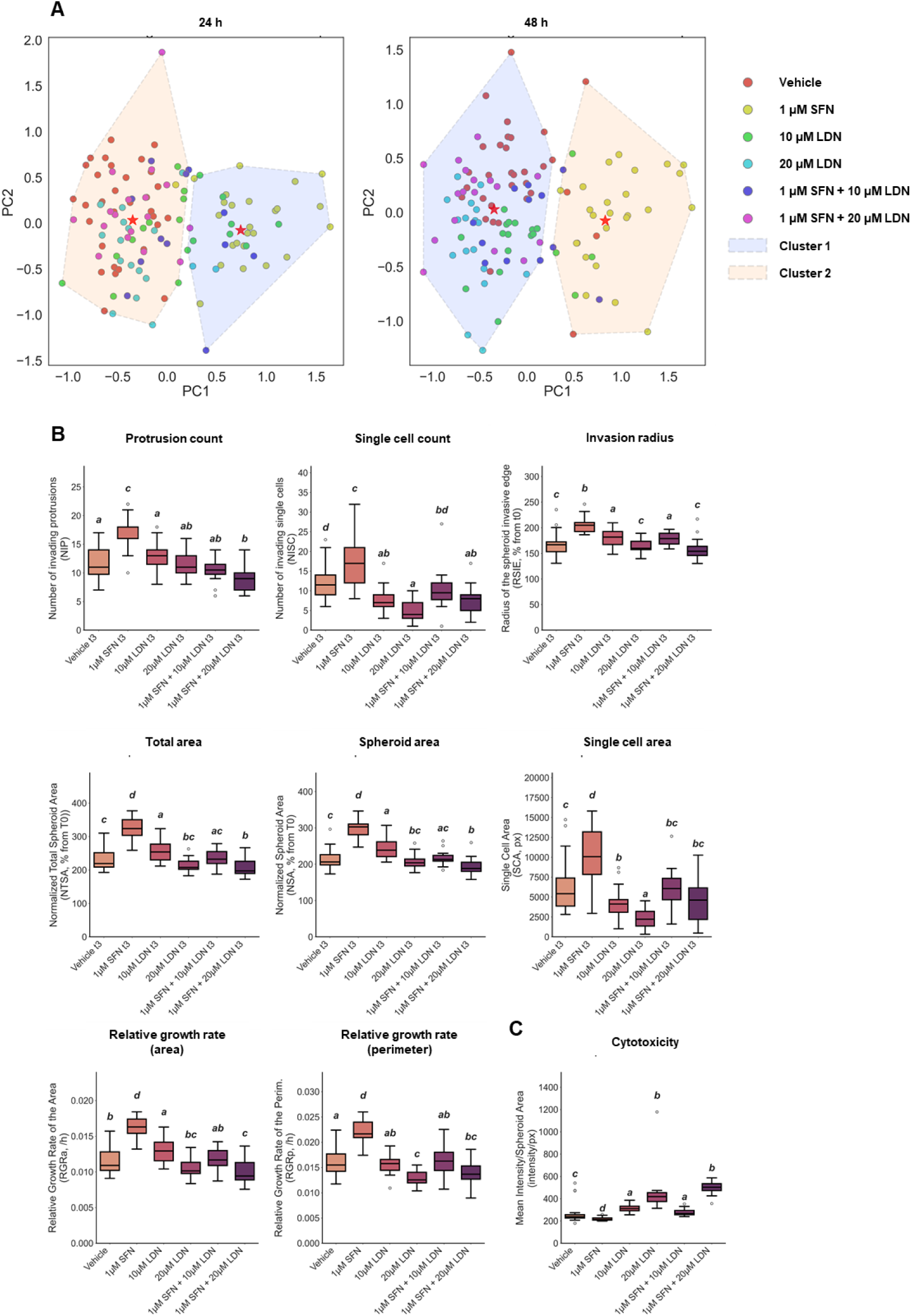
Co-treatment with SFN and LDN reverses the SFN-induced invasion shift in Hep3B spheroids. (A) Multiparametric clustering (PCA + k-means) after 24 h and 48h treatment indicating co-treated spheroids cluster with the vehicle and mono-LDN rather than with SFN 1 µM. This indicates that the SFN-induced invasion shift is countered when combined with LDN treatment. Dots represent spheroids and n/condition are for DMSO: 32, for 1µM SFN: 25, for 10µM LDN: 19, for 20µM LDN: 17, for 1µM SFN + 10µM LDN: 12, for 1µM SFN + 20µM LDN: 17. PCA is based on an extensive set of growth, invasion, morphology and spheroid complexity features. (B) Boxplots (median and interquartile range (IQR)) for selected relevant invasion and growth features. Features shown describe the reversal of invasiveness upon co-treatment with SFN and LDN after 72h treatment. Included are: Number of protrusions (counted as the number of protrusions on the edge of the main spheroid area), the number of invading single cells/areas, the invasion radius (defined as the radius of the outer bounding circle i.e. the smallest circle encompassing the spheroid and all invading areas), the total area (which includes the summed area of the central spheroid mass (spheroid area) and all invading areas (single cell area), the relative growth rate (RGR) of the area and perimeter (see Fig. S3). (C) Cytotoxicity of spheroids based on mean intensity of SYTOX green staining within the spheroid boundary (median and interquartile range (IQR)). Increased toxicity is not significant for the combinations beyond the toxicity of the mono-treatments. Toxicity levels are in general lower than e.g. for high SFN (see 10 µL in Fig. S3D). (B-C) Statistical significance is shown as compact letter display: letters (a, b, c) indicate non-significant groups (p > 0.05) (ANOVA/Tukey or Kruskal–Wallis/Dunn depending on normality).

**Fig. S6.**
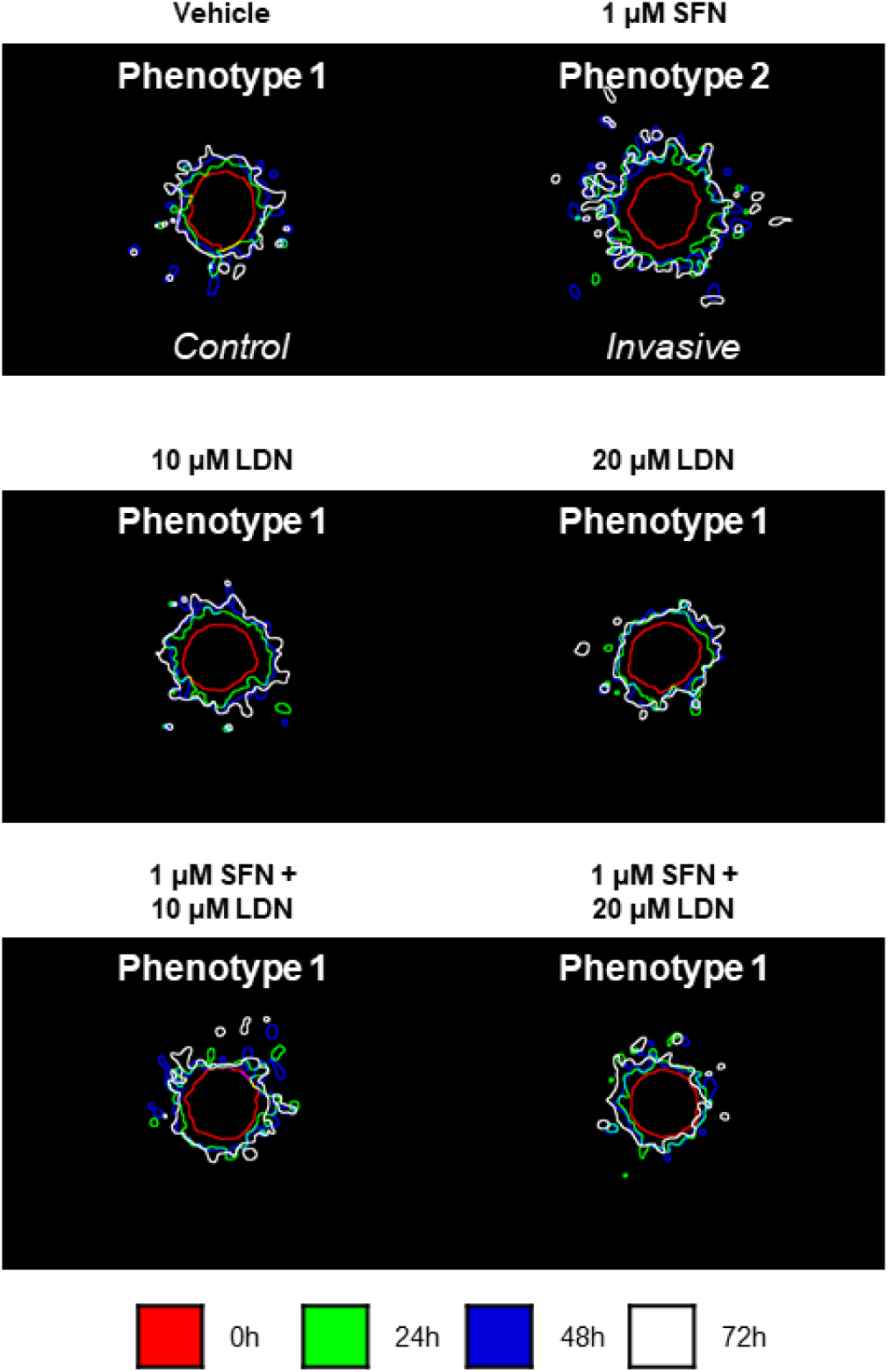
Phenotypes of Hep3B spheroids at different treatment times with SFN, LDN or combinations. Overlay of spheroid outlines in time generated by SImBA software (ref). The outlines at different time points are shown in different colors (see legend). Two phenotypes were distinguished based on the analysis shown in Figure S3: phenotype 1 (control condition) and phenotype 2 (invasive condition for 1µM SFN). Based on the Figure S3 analysis, these phenotypes are here assigned to the overlays of the representative spheroids of each treatment condition. Vehicle indicates control condition (DMSO). This demonstrates that LDN has no effect on its own (phenotype 1) but in combination with SFN reverts the SFN-induced pro-invasive phenotype 2 back to phenotype 1.

**Fig. S7.**
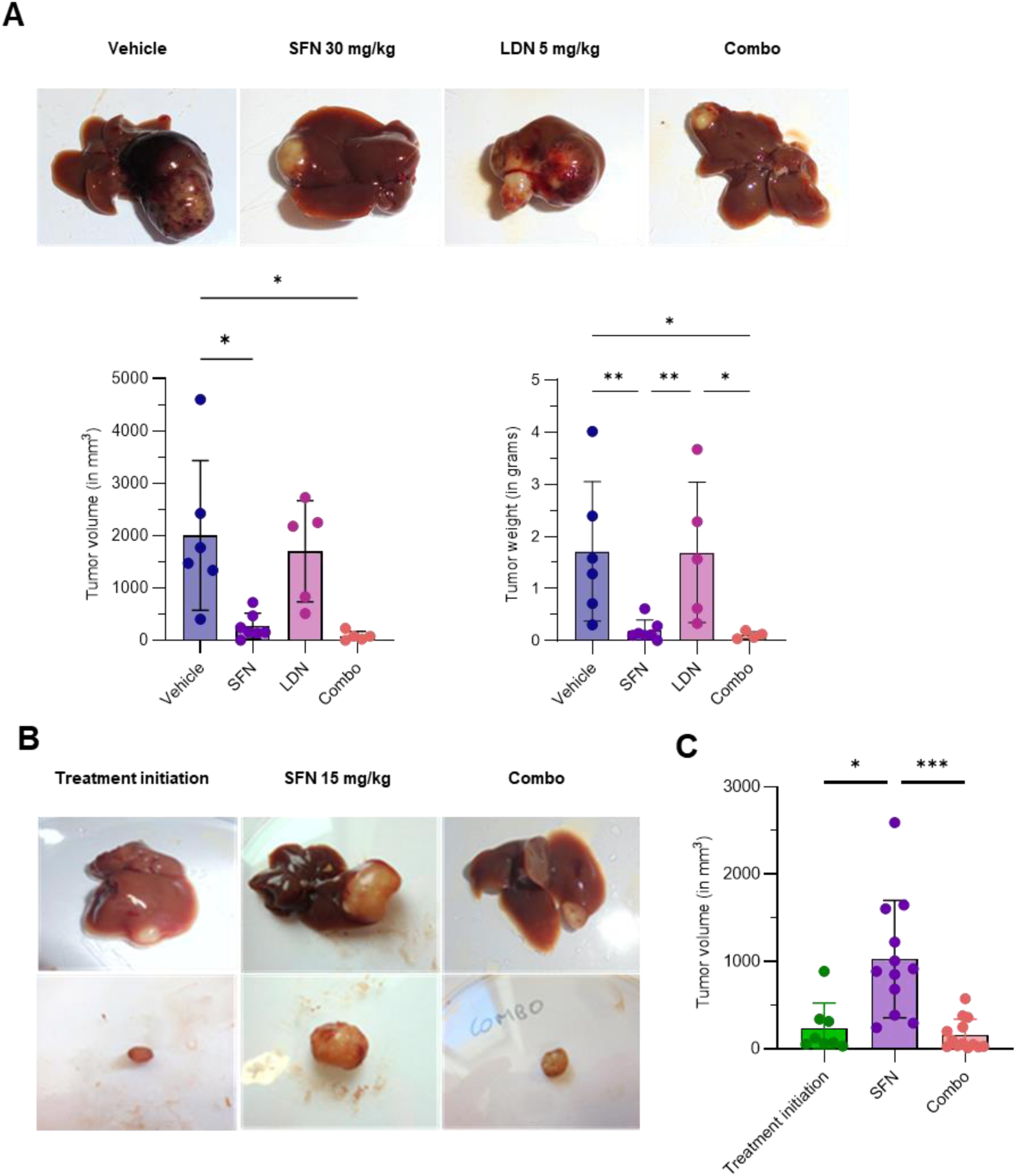
SFN-LDN co-treatment effects on tumour growth in orthotopic mouse model at different SFN doses. (A) Representative tumour images (top) and quantification of tumour volume (bottom left) and tumour weight (bottom right) from Hep3B xenografts treated with vehicle, higher SFN (30 mg/kg), LDN (5 mg/kg), or their combination. SFN monotherapy already produced a pronounced reduction in tumour burden versus vehicle, indicating that this SFN dose is already highly effective in vivo. The combination, although resulting in the lowest absolute tumour burden, only showed a modest, nonsignificant benefit over SFN alone. Data are mean ± SD; *:p<0.05, ** p<0.01 based on one-way ANOVA with post-hoc testing. (B) Representative macroscopic images of Hep3B xenograft tumours at treatment initiation (baseline), after SFN monotherapy (15 mg/kg), or after SFN–LDN combination treatment (15 mg/kg SFN, 5 mg/kg LDN). Images show both liver-embedded tumours (top) and excised tumour masses (bottom). (C) Quantification of tumour volume at the start of treatment and after three weeks treatment as in B: SFN (15 mg/kg) or SFN–LDN therapy (15 mg/kg SFN, 5 mg/kg LDN), demonstrating not only indicating superiority over SFN monotherapy at this low SFN dose but also near-complete suppression of tumour growth. Data are mean ± SD; *p<0.05, **: p<0.01, ***p<0.001 based on one-way ANOVA with post-hoc testing.

